# Interspecies systems biology links bacterial metabolic pathways to nematode gene expression, behavior, and survival

**DOI:** 10.1101/2024.10.02.616249

**Authors:** Marina Athanasouli, Tobias Loschko, Christian Rödelsperger

## Abstract

All animals live in tight association with complex microbial communities, yet studying the effects of individual bacteria remains challenging. Here, we exploit the system of the bacterial feeding nematode *Pristionchus pacificus*, where specific interactions can be studied in isolation. We sequenced the genomes of 84 *Pristionchus*-associated bacteria to investigate how differences in bacterial metabolic potential affect nematode traits. Specifically, we identified the Paerucumarin biosynthesis pathway as a strong candidate for nematode survival. Hypothesizing that bacterial production of coumarin derivatives contributed to increased lethality, we demonstrated nematicidal effects of two related compounds using supplementation assays. To explore metabolic interactions at a broader scale, we generated nematode transcriptomes for 38 bacterial diets and constructed a coexpression network that revealed distinct developmental and environmental signatures. Interspecies association studies resulted in a bipartite network with more than 2,800 interactions which may assist future studies to identify metabolites affecting various biological processes.

## Introduction

The human gut microbiome exemplifies the intricate associations between organisms and bacteria, highlighting the complexity of cross-kingdom interactions and their impact on host development and disease. These interactions are influenced by the heterogeneity of the gut microbiome, as well as variations in host diet and genetic background, making them challenging to study. In recent years, the nematode *Caenorhabditis elegans* has emerged as a powerful model for investigating host–microbiota interactions. This model allows for detailed examination of individual interactions through the use of monoxenic bacterial cultures, where worms are grown on single bacterial strains^1^. *C. elegans* engages with bacteria in multiple ways: it is a bacterial feeder, typically consuming the *Escherichia coli* OP50 strain in laboratory settings, and the bacterial diet significantly influences its development, as exemplified by the vitamin B_12_ production by *Comamonas* that impacts *C. elegans* development^2^. Moreover, *C. elegans* harbors a complex microbiome in its natural habitat, with certain bacteria enhancing the host’s fitness under stress^3, 4^, such as *Bacillus subtilis*, which provides resistance against Gram-positive pathogens when colonizing the nematode’s gut^5^.

The nematode *Pristionchus pacificus* was initially introduced as a satellite model organism for comparative studies with *C. elegans*. Having diverged approximately 130-310 million years ago^6^, these species share several traits, including transparency, short generation times, and hermaphroditism, which make them valuable models for genetic research. More recently, *P. pacificus* has gained prominence as a model for studying the interplay between environment and genotype in producing diverse phenotypes. A key example of this is the species’ mouthform plasticity, where genetically identical individuals can develop either a narrow mouth suited for bacterial feeding or a wider mouth with an additional tooth for predation. This plasticity, regulated both genetically and environmentally, varies among natural isolates and is influenced by dietary factors. For instance, the reference *P. pacificus* strain PS312 predominantly produces the predatory morph when fed *E. coli* OP50 or other bacteria but shifts to the non-predatory morph when consuming a specific *Cryptococcus* yeast strain^7^. Additionally, factors such as specific pheromones and alternative culture conditions have been shown to affect mouthform plasticity^8, 9^. Despite the extensive characterization of the regulatory network controlling this developmental switch^10-14^, little is known about how *P. pacificus* generally processes environmental signals. Our focus is on the nematode’s interactions with environmental microbiota, as *P. pacificus* is often found in decaying scarab beetles, where it feeds on various microbes^15, 16^. Notably, one major difference between *C. elegans* and *P. pacificus* is that the latter lacks a specific morphological structure, the grinder, that is used by *C. elegans* to physically lyse the bacteria. This likely has widespread consequences on the way *P. pacificus* interacts with bacteria including pathogen susceptibility^17, 18^. Previously, we isolated and cultured 136 bacterial strains from *Pristionchus*-associated environments and demonstrated that *P. pacificus* exhibits differential development, behavior, and survival in response to these bacteria^19, 20^. Notably, we found that bacteria-derived vitamin B_12_ not only accelerates development but also enhances predatory behavior.

In this study, we apply a systems biology approach to investigate the interactions between *Pristionchus pacificus* and bacterial food sources derived from the nematode’s natural environment. Specifically, based on whole-genome sequencing, we explore the variation in metabolic potential among these bacteria and we test if this variation can be associated with the differential responses of the nematodes when exposed to these bacteria. Our main goal is to identify potential interactions between bacterial metabolic pathways and nematode phenotypes which include previously generated data on survival and chemotaxis behavior^19^. We perform supplementation experiments to further support that a specific group of metabolites could potentially contribute to the high toxicity of some bacteria. Further, to investigate metabolic interactions at a broader scale, we generated transcriptomic profiles of worms grown on 38 different bacteria and tested for associations between *P. pacificus* coexpression modules and bacterial metabolic pathways. This resulted in a large bipartite network involving more than 2,800 interactions, which highlights the complexity of the whole system and serves as a resource for future studies elucidating how specific metabolites control developmental and metabolic processes in *P. pacificus*.

## Results

### Sequencing of 84 bacteria establishes the genomic basis to study host microbe interactions

In order to link genomic variation in bacteria to differential responses in *P. pacificus* nematodes, we sequenced 84 strains from a large bacterial collection isolated previously from *Pristionchus*-associated environments^19^. This collection mostly comprised Proteobacteria as well as some members of the phyla Firmicutes, Actinobacteria, and Bacteroidetes. Among the Proteobacteria, the most deeply sampled families constitute Enterobacteriaceae, Pseudomonadaceae, and Moraxellaceae. 84 bacterial strains were selected for whole genome sequencing based on criteria such as fast growth, easiness of DNA extraction, low propensity of contamination. (Supplemental Table S1). The resulting genome assemblies were highly complete as indicated by a median single-copy BUSCO completeness of 99% with an interquartile range (IQR) of 98-100%. The largest parts of the bacterial genomes could be assembled into contigs spanning more than 100kb. Specifically, the N50 value, which indicates the minimum contig length in the set of largest contigs that account for at least half of the genome, was 324kb (IQR=107-498kb) and median genome assembly size was 4.9Mb (IQR=4.4-5.6Mb). Gene annotation yielded a median of 4,562 gene models per assembly (IQR=4,184-5,225) with a median single-copy BUSCO completeness level of 99% (IQR=98-100%). The complete overview about these different quality measures can be found in Supplemental Table S1. This set of bacterial genomes builds the basis to study host microbe interaction in the *Pristionchus* system.

### *Pristionchus*-associated microbiota harbor previously uncharacterized bacterial strains

The predicted proteomes of all 84 genomes were taken to reconstruct a bacterial phylogeny (see Methods) (Fig. 1). The different lineages in the resulting species tree were mostly consistent with previous genus assignments based on 16S sequencing^19^. To more accurately classify the strains, we performed homology searches against the non-redundant version of NCBI and determined the nomenclature of our bacteria by assigning each strain to the bacterial genus with the majority of best hits (Supplemental Table S2). During this process, we observed that the median percentage of identity of the strains *Sphingobacterium* L2, *Pseudomonas* L74 and *Erwinia* V71 to their respective best hits in NCBI was lower than 90%. Additionally, we were unable to definitively determine the genus for the closely related strains V69 and V91, despite a median identity of 90.6% and 96% respectively. We hypothesized that these bacterial strains could either represent novel isolates or have no associated whole genome sequencing data in the NCBI database. To confirm our hypothesis regarding the potentially novel bacteria, we searched the literature for recent, comprehensive phylogenies of the genera^21-24^ and recreated these phylogenies including our strains and their best hits in the NCBI database. The reconstructed phylogenies for the three bacterial strains with low median identity (L2, L74, V71) revealed that our strains are outgroups to the clusters containing their best hits in their respective phylogenetic trees but were still more closely related to them compared to any other genus (Supplemental Fig. 1A-C). The reconstructed phylogeny including V69 and V91 exhibited the same patterns and indicated that both bacteria should be classified as *Phytobacter* strains (Supplemental Fig. 1D). These findings support that *Pristionchus*-associated microbiota constitute a previously undersampled environment that harbors novel bacterial strains and species.

**Figure 1.**
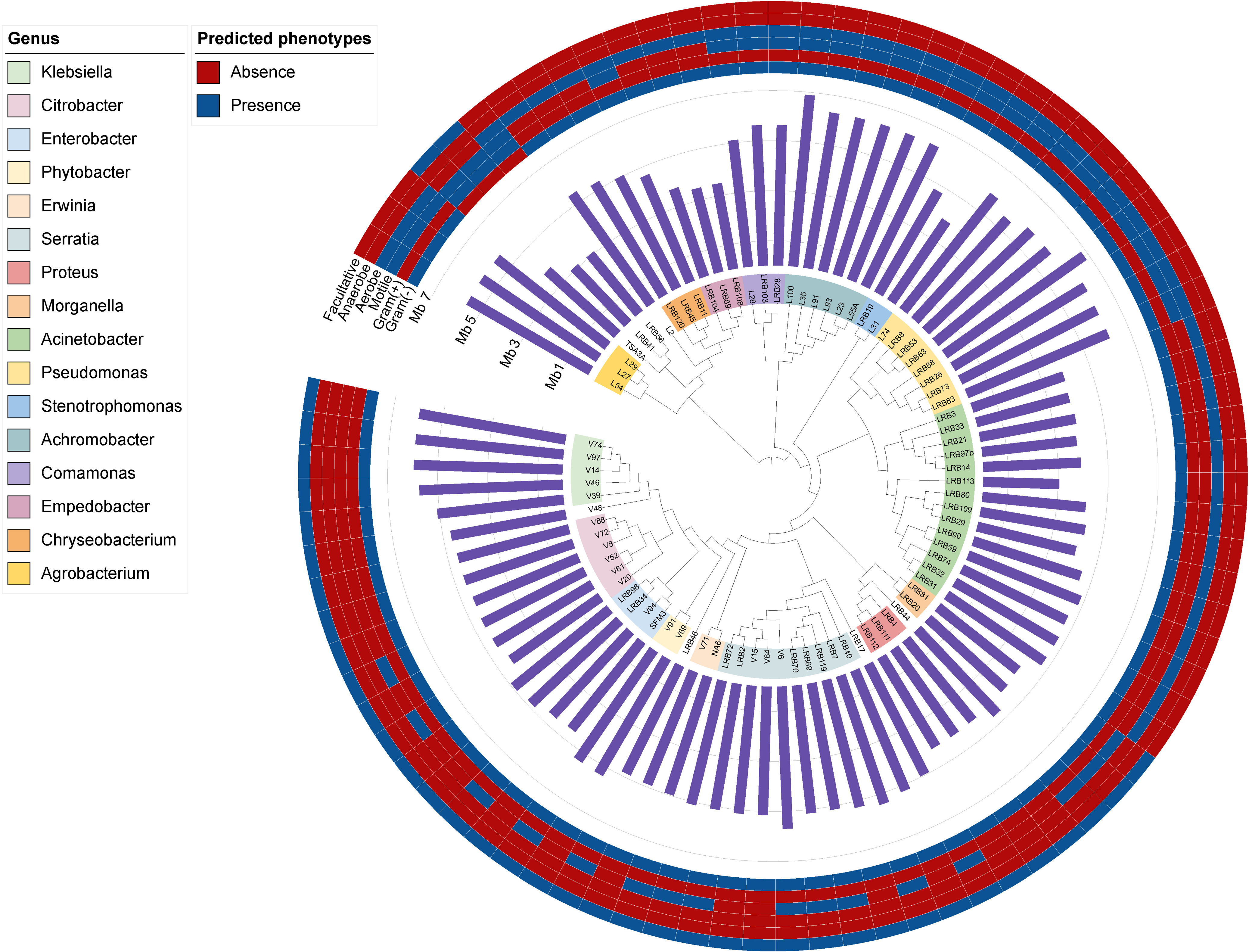
Phylogenomics of 84 bacterial genomes. The tree depicts the bacterial phylogeny acquired using RAxML after whole genome sequencing, genome assembly and orthogroup detection. Phenotype prediction with traitar3 revealed that the overwhelming majority of bacterial strains sequenced are anaerobic with the exception of TSA3A and LRB41 who are facultative anaerobic as well as the complete lack of anaerobes. Our collection contains only three Gram positive strains while the rest are gram negative. Genome sizes may vary even within a genus as is the case with *Acinetobacter*, where we observe genomes ranging between 3 and 5 Mb.

### Closely related bacteria exhibit substantial variation in metabolic potential

In order to characterize the variation of metabolic potential among members of the *Pristionchus*-associated microbiota, we reconstructed metabolic networks for the 84 bacterial genomes with the gapseq approach using existing pathway information from the MetaCyc database. MetaCyc was preferred to KEGG due to the higher detail it offers with regards to the available pathways. We then investigated the absence/presence patterns of metabolic pathways among the 84 bacteria and defined metabolic pathways groups (MPGs) based on identical patterns (Fig. 2A). This reduced the number of total metabolic pathways from 2,902 (Supplemental Table S3) to 715 MPGs. The two most abundant MPGs denote 1,832 and 45 individual pathways that are either absent or present in all strains (Fig. 2A). While most pathways that were not detected in our data correspond to pathways that are known only from eukaryotes, examples of the core metabolic pathways that are present in all strains relate to the TCA cycle, biosynthesis of several amino acids as well as fatty acid biosynthesis and elongation pathways, pointing towards necessary pathways for the survival of bacterial cells (Supplemental Table S2). The third most abundant MPG 3 denotes 13 pathways that are only missing in *Enterococcus* strain TSA3A which is one of only three Gram positive strains in our collection (Fig. 1). Its genome assembly has good contiguity (N50=234kb) and is highly complete (BUSCO completeness = 98%), which suggests that this pattern is unlikely due to a lower quality of this particular genome. Also, more detailed analysis of these 13 candidate pathways revealed that while some pathways are partially present other pathways are completely absent. Other MPGs show strong phylogenetic signatures such as MPG 7 and 13 that are restricted to the genera *Achromobacter* and *Pseudomonas*, respectively. We would therefore conclude that most of the variation in metabolic potential rather reflects genome evolution than technical issues. In order to better quantify the metabolic variation in our bacterial collection, we counted the number of pathway differences for pairwise comparison within and across genera. This showed significantly fewer differences in the metabolic potential for strains of the same genus (Fig. 2B). However, even strains of the same genus can differ in dozens of metabolic pathways, which can potentially explain the previously observed variability in the molecular and phenotypic responses of *P. pacificus* nematodes^19, 25^.

**Figure 2.**
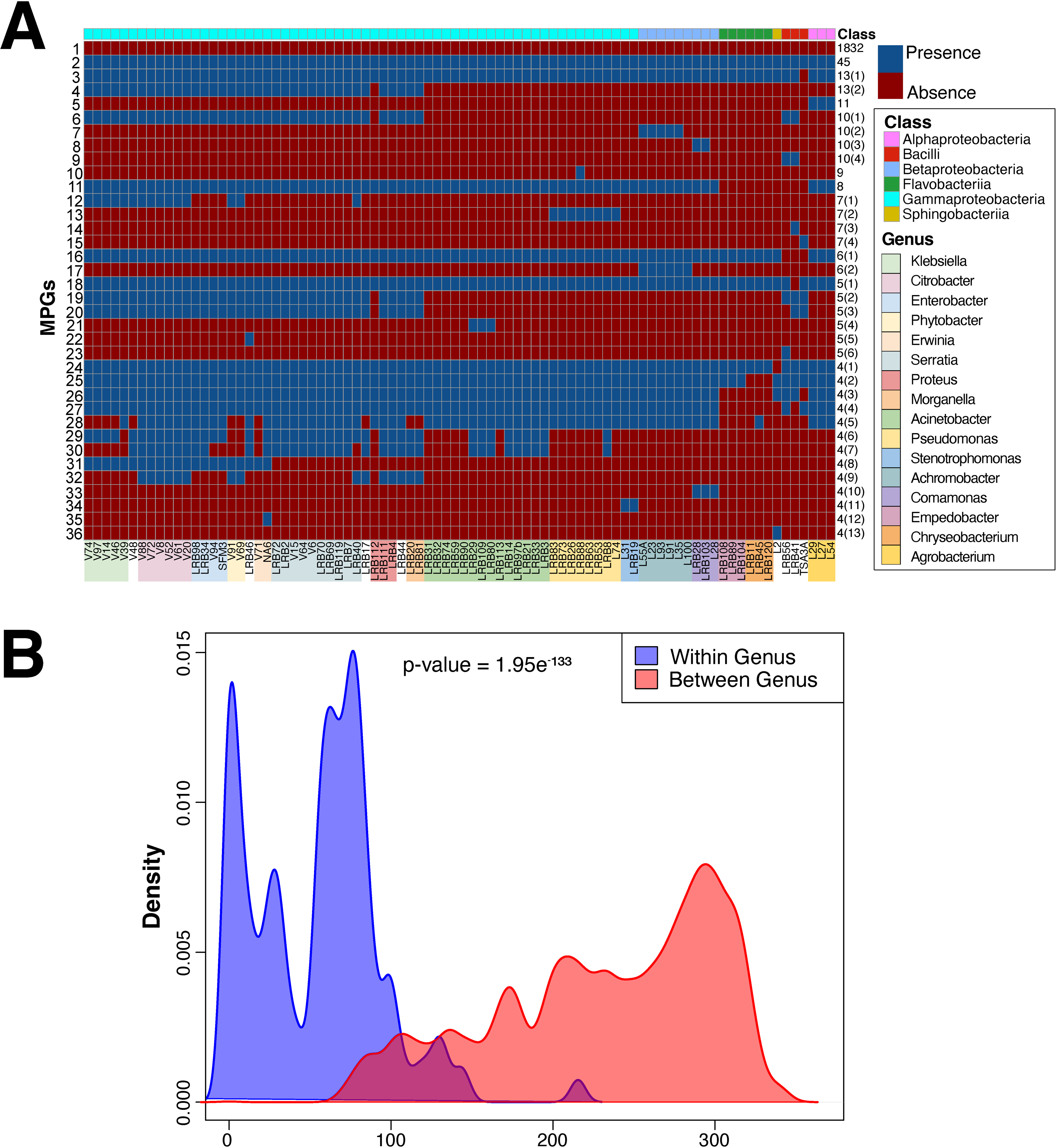
Variation in bacterial metabolic potential. (A) Grouping of the predicted MPGs depending on their presence or absence across the bacterial phylogeny revealed the metabolic potential of the individual strains. The majority of MPGs detected were absent from all strains with a core of 45 bacterial pathways predicted always present. The rest of the MPGs exhibit distinct patterns even among the same genus. (B) Pairwise comparisons of the MPGs present in strains of the same genus versus strains of different genera indicated that closely related bacteria share a higher number of MPGs and thus, might exhibit a genus-specific metabolic signature.

### Interspecies association studies identify candidate pathways affecting nematode survival and behavior

During the initial culture-based characterization of *Pristionchus-*associated microbiota, Akduman et. al. performed assays for chemotaxis behavior and survival with over a hundred bacterial strains^19^, many of which are part of our collection (Fig. 3A). In a pilot experiment, we employed our genomic data to screen for candidate pathways, and their associated metabolic products, that could possibly impact nematode survival and behavior. To this end, we tested for associations between the presence or absence of a MPG with the behavioral and survival phenotypic data mentioned above. With this strategy, we identified 24 MPGs exhibiting significant association with chemotaxis (*P* < 0.05, Wilcoxon-test, Supplemental Fig. S2 and Table S4) and 40 MPGs that are associated with survival (Fig. 3B, Supplemental Table S5). Among these candidate pathways, we noticed a dramatic difference in survival associated with between presence or absence of the paerucumarin and rhabduscin biosynthesis pathway. Coumarin-derivatives and rhabduscin have frequently been reported to exhibit nematicidal activity^26-29^, which suggests that similar bacteria-derived metabolites contribute to the reduced survival of *P. pacificus* nematodes. To complement this analysis with an additional screen for other virulence factors, we combined the predicted proteomes of the 84 bacteria with the virulence factor database (VFDB)^30^ and performed a similar association study. This yielded 15 orthologous clusters with significant association (FDR-corrected P-value < 0.05), among which members of the paerucumarin biosynthesis gene cluster showed the strongest and most consistent effect (Supplemental Fig. S3). Closer examination of the presence of these genes in our data set revealed that these orthologs are exclusively present in strains of the genus *Serratia.* Since the paerucumarin gene cluster has been initially characterized in *Pseudomonas*^31^, the homologs of those genes in *Serratia* will not necessarily produce the exactly same compound. Thus, hypothesizing that paerucumarin, rhabduscin, or similar metabolites could contribute to the pathogenicity of *Serratia* strains, we tested three coumarin-derivatives, 4-Hydroxycoumarin, 7-Methoxycoumarin, and Xanthotoxin, that were previously shown to possess some nematicidal activity and that are commercially available for their effect against *P. pacificus*^27-29^. Among those, 7-Methoxy coumarin and Xanthotoxin caused substantial lethality at 1mM concentrations (P < 0.05, t-test, Fig. 3C). This supports that some coumarin derivatives have nematicidal activity against *P. pacificus* and that such compounds could potentially contribute to the increased pathogenicity of *Serratia* strains. However, further experimental work will be needed to identify which metabolites are produced by the *Serratia* strains and whether they exhibit indeed nematicidal activity.

**Figure 3.**
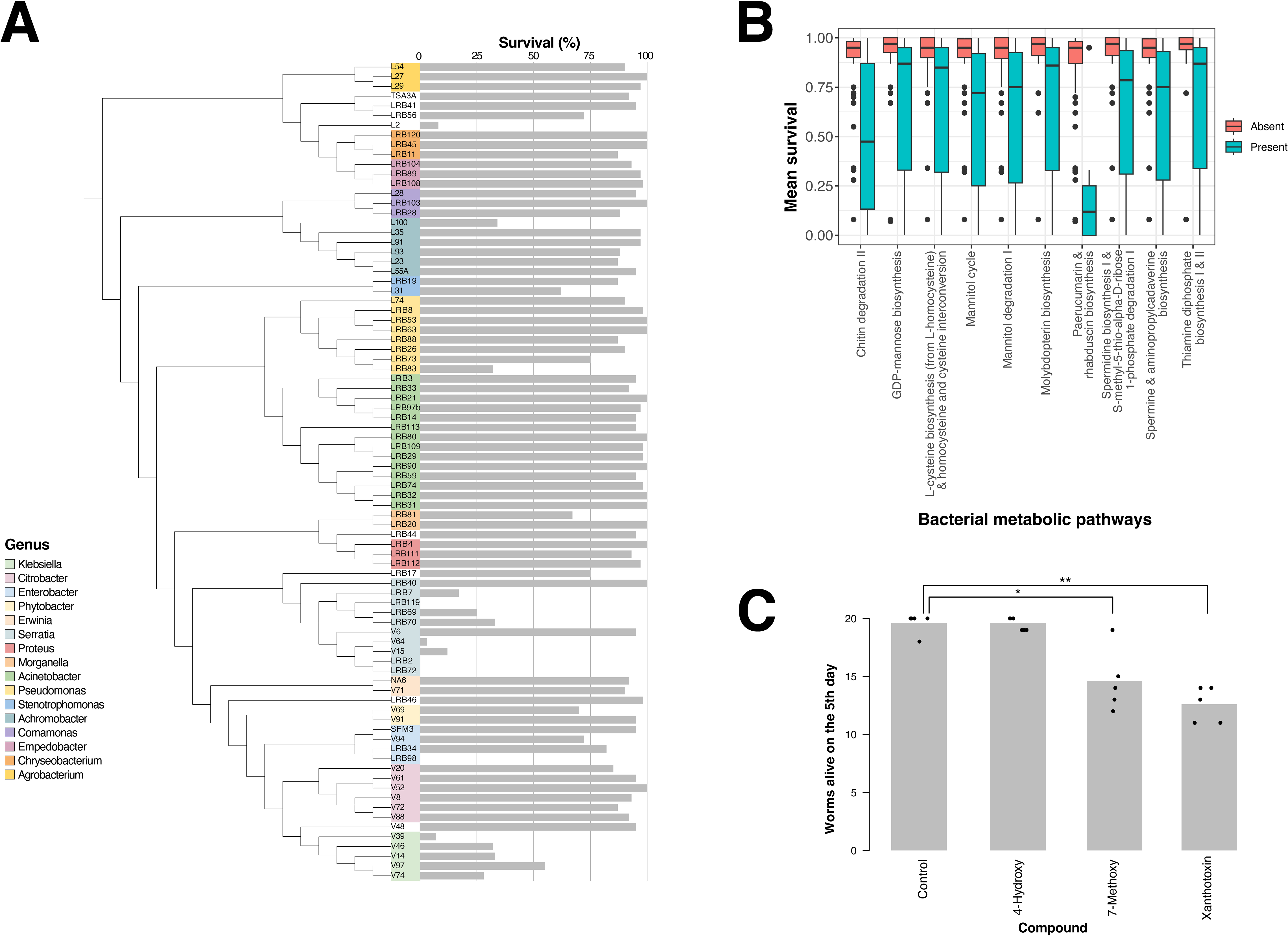
Association of nematode survival with the bacterial pathways. (A) An association study between the MPGs and mean survival data narrowed down the MPGs with a significant effect on the trait from hundreds to 41, a subsample of which is shown here. Generally, the presence of an MPG appears to result in lower survival, especially when bacteria produce chitinases and compounds such as paerucumarin and rhabduscin. (B) Supplementation tests with three types of coumarins showed that 7-methoxy coumarin and xanthotoxin affect nematode survival (P<0.01, t-test).

### Closely related bacteria can induce vastly different transcriptomic responses in nematodes

Having demonstrated the utility of the bacterial genomes to uncover candidate pathways impacting nematode survival and behavior, we now focus on our main objective to identify interactions between bacterial metabolic pathways and regulatory and metabolic response networks in the worm. To this end, we generated nematode transcriptome profiles for a subset of 38 dietary bacteria that belong to the three classes, Alpha-, Beta- and Gammaproteobacteria, and represent 16 genera. These 38 bacteria were selected as they supported complete development from egg to adult for at least two generations, which was not the case for all the strains. For each bacterial strain, we generated two biological replicates by collecting F1 worms 72 hours after egg-laying (see *Methods*). While most animals should have reached adulthood at this time under standard *E. coli* OP50 diet^32^, we cannot exclude that some bacterial diets either accelerate or slow down development leading to a shift of the transcriptomic profiles towards either older or younger developmental stages. We decided to sequence the transcriptomes of mixed-stage populations as we observed previously that even when adult worms are manually picked from plates, a developmental signature is still visible in the resulting RNA-seq data. This is because worms will still develop at different speeds and chronological age might not necessarily correspond to developmental age^25^. As a consequence, differences in stage composition likely explain the larger extent of transcriptomic variation in our study, i.e. quite a number of samples showed correlation coefficients r < 0.9 (Fig. 4A), which was not the case in our previous study where we manually picked adult worms on 24 diverse bacteria^25^. We note that biological replicates do not always group together suggesting that the effect of experimental variation on the overall transcriptome can in individual cases be larger than the effect of different bacterial diets. However, generally, we observed that replicates of the same bacterial diet exhibit less variation and thus are more similar than transcriptomic profiles from different bacterial diets (Fig. 4B). With regard to the taxonomic grouping of the bacteria, we observe that bacterial strains of the same genus show more similar transcriptomic responses in the worms than members of different genera (Fig. 4C). However, even in comparisons within the same genus, transcriptomic responses can differ substantially with correlation coefficients < 0.9. This recapitulates our previous finding of substantial variation in bacteria from the same genus with regard to metabolic potential (Fig. 2B).

**Figure 4.**
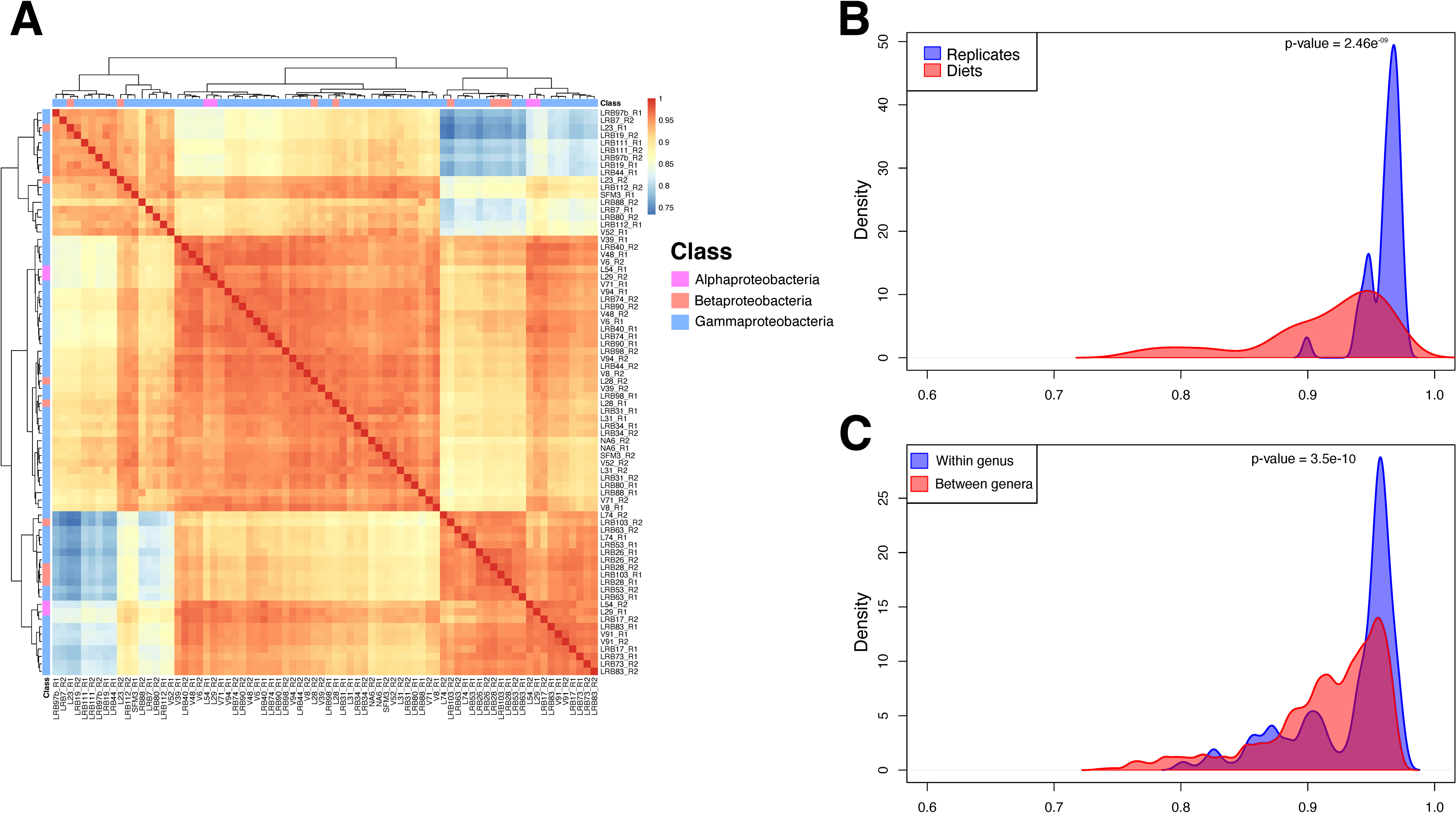
Variation in the transcriptomic response of the nematode. (A) Correlation analysis and clustering of the transcriptomes from *P. pacificus* nematodes in response to 38 different bacteria. The overall transcriptomes are highly similar with most pairwise comparisons showing correlation coefficients >0.9. (B) Biological replicates exhibited higher correlation coefficients compared to transcriptomes from different bacterial diets. (C) Transcriptomes of bacteria from the same genus showed significantly higher correlation coefficients compared to dietary samples from different genera.

### Specific network modules respond to signals from diverse bacteria

While the above mentioned transcriptomic analysis captures overall transcriptomic similarities, specific effects of individual bacteria might be overshadowed by global patterns. To investigate the effects of individual bacteria on smaller gene sets that may point towards specific regulatory or metabolic network modules, we constructed a gene coexpression network from the RNA-seq data following the protocol and parameters from our previous related work^25^. Overall, our coexpression network captures 24,375 (84%) of *P. pacificus* genes, of which 13,972 are found in the 60 largest modules with more than 20 genes. Visualization of expression values in these modules across the different microbiota demonstrated that individual modules appear to be induced by specific bacteria (Fig. 5). For example, the second largest module 2 shows high expression on most *Pseudomonas* strains whereas module 7 exhibits highest expression only on the *Pseudomas protegens* strain LRB88. Note that some of these patterns may represent indirect effects of altered development on diverse bacterial diets. This is shown by distinct expression signatures of specific coexpression modules when overlaying them with the developmental transcriptomic data of *P. pacificus* (Supplemental Fig. 4)^32^. For example, module 1 peaks late in development close to adulthood and is slightly preceded by a peak of module 2. However, no matter if the expression of those genes is a direct or indirect consequence, it can be interpreted as a specific response of *P. pacificus* nematodes to a distinct microbiota^25^.

**Figure 5.**
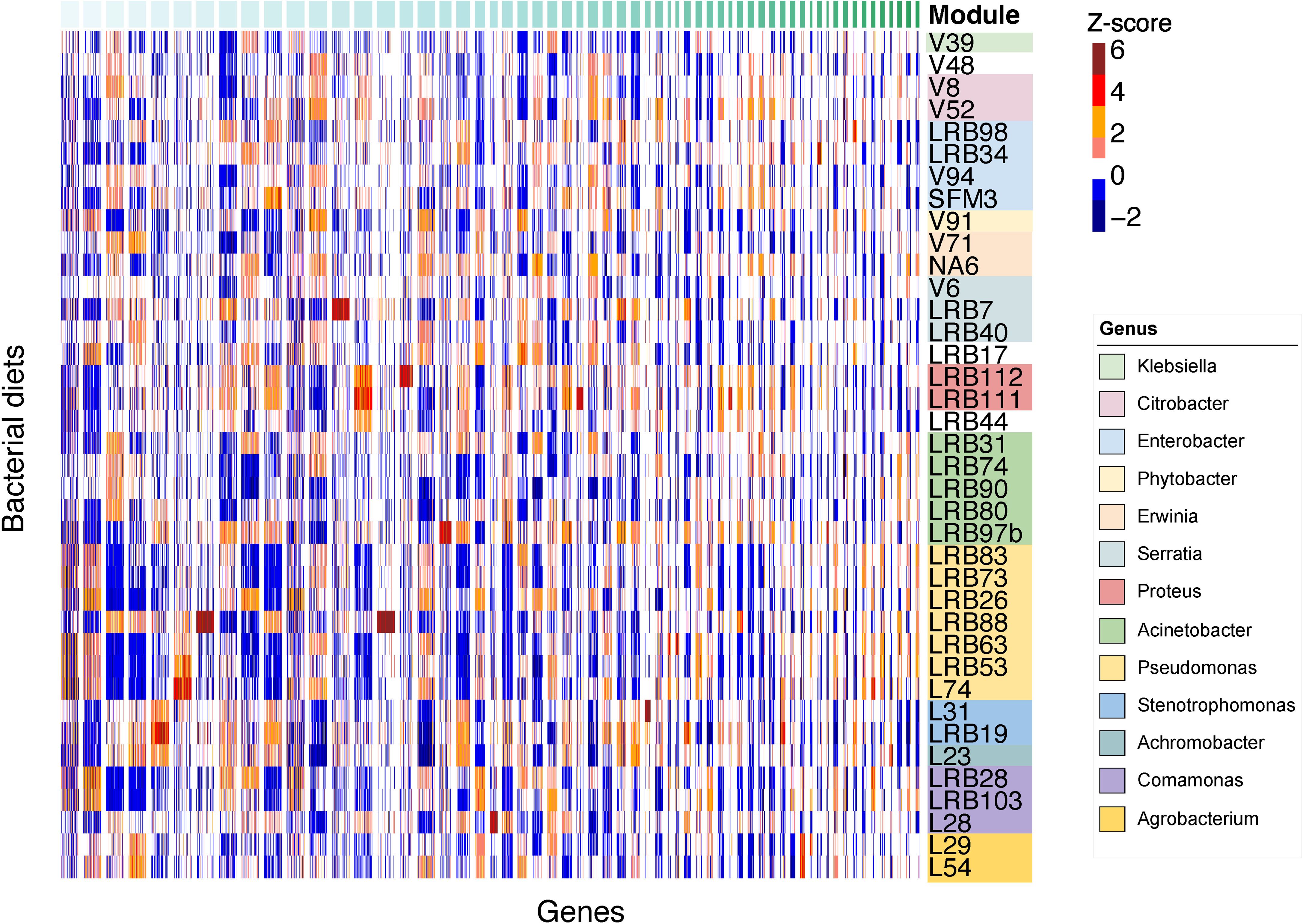
Expression of modules across the different bacterial diets. The z-score normalized expression of the top 60 coexpression modules across the different diets was visualized as a heatmap. The expression of the biological replicates was averaged in order to produce a single expression profile for each of the 38 diets. Modules with more than 50 genes were randomly downsampled to aid the visualization of smaller modules and the comparisons between them. Expression patterns were unique for each module and depended on the bacterial diet, as was the case with modules 2, 7 and 15 on LRB88.

### Most coexpression modules can be functionally annotated

To functionally annotate the coexpression modules and to gain insight into the regulation of metabolic pathways, we analyzed the top 60 modules of the network that contained more than 20 genes and performed enrichment analyses, similar to previous work. This included overrepresentation of protein domains^33^, regionally enriched genes as identified from spatial transcriptomics^34^, KEGG pathways^35^, and previous gene expression studies focusing on sex-biased genes^36^, intestinal transcriptome^18^, and developmental oscillations^32^ (Supplemental Table S6). This allowed the labeling of 48 modules with biological terms based on the strong enrichment patterns (Fig. 6, Supplemental Table S7). For example, the largest module 1 (OOGEN_1) is overrepresented in hermaphrodite-biased and gonad-enriched genes (Fig 6). It also includes known germline expressed markers like *spo-11*, *dmc-1*, *hcp-4* (*cenp-c*), *syp-4*, and *hop-1*^37^, indicating that it largely represents oogenesis. However, it also shows some overrepresentation of neuropeptides and head-enriched genes suggesting that also a portion of neuronal genes are captured in this module. The second largest module 2 (SPERM_2) seems to be more homogenous, as all overrepresented terms point towards spermatogenesis (Fig. 6). The assignments of module 1 and 2 are also in line with their developmental signature (Supplemental Fig. 4) as sperm are generated during late larval development whereas oocytes are produced during adulthood. While many modules show less specific signals, individual modules show specific enrichments with individual pathways or anatomical regions. For example, module 36 (GLAND_36) is strongly associated with gland cell specific expression^13^. We used the microbial transcriptomes (Fig. 5), to label a handful of modules that did not show any enrichment in the above mentioned data set (Supplemental Table S7). Thus, module 13 (LRB7_13) is strongly upregulated on *Serratia quinivorans* strain LRB7 and module 27 (LRB111_27) exhibits highest expression on *Proteus terrae* strain LRB111. Altogether, the characterization of the 60 largest coexpression modules functionally annotates a large portion of the *P. pacificus* gene set which includes many lineage-specific genes for which no homologs are known in *C. elegans*^16, 25^.

**Figure 6.**
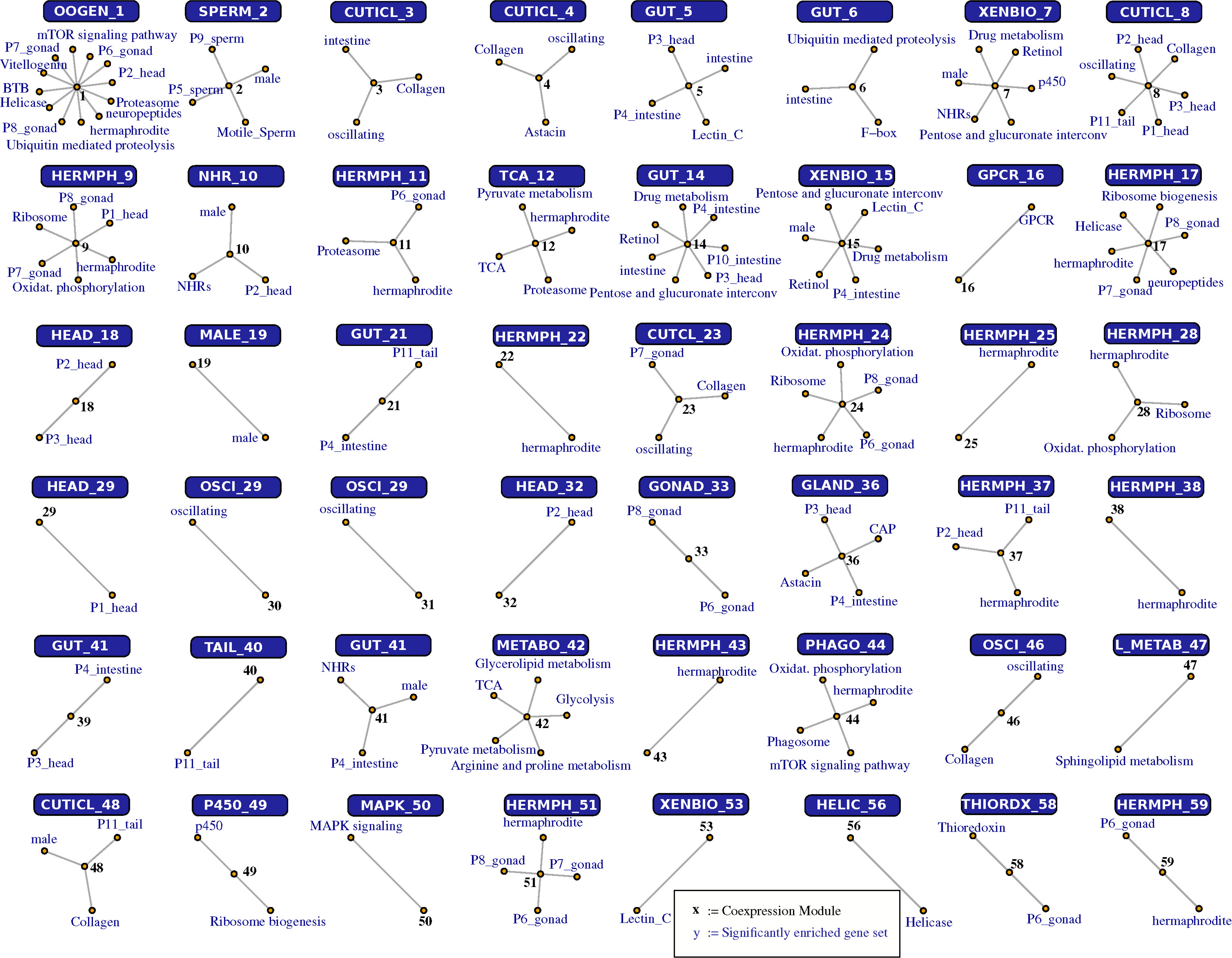
Enrichment analysis of the gene coexpression modules using existing transcriptomic datasets and functional annotation. The networks show selected results of the overrepresentation analysis for specific coexpression modules. See Supplemental Table S6 for the full data set. Based on the strongest enrichment patterns, we connected 48 of the processed 60 modules with biological processes and protein domains, allowing the functional labeling of the majority of the modules (Supplemental Table S7).

### Interspecies association studies traces the response of coexpression modules to specific bacterial pathways

A main goal of this study was to trace back the transcriptomic response of individual coexpression modules in *P. pacificus* to candidate metabolic pathways in the bacteria. We hypothesized that some of the MPGs show strong associations with our coexpression modules. To this end, we combined the bacterial MPGs and the nematode coexpression modules into a bipartite network and calculated the interaction of each MPG with the modules based on the significance of the overlap between the genes within a coexpression module and the genes that exhibit differential expression in response to the absence or presence of an MPG. Altogether, we observed more than 2,800 interactions that involved 620 MPGs and 42 coexpression modules (Supplemental Table S8). The module with the highest number of predicted interactions was intestinal module 14 (GUT_14). Furthermore, the majority of coexpression modules interacting significantly with at least 100 MPGs are intestinal, a finding highlighting the important role of the intestine and is consistent with recent studies^38^. To visualize the strongest interactions in our enriched dataset, we chose the 50 MPGs with the lowest Bonferroni-corrected p-values (Fig. 7). Surprisingly, the largest module of the network (OOGEN_1) exhibited interactions with only 46 MPGs (Supplemental Table S8), of which 21 are highly significant (Fig. 7). On the contrary, module 2 (SPERM_2) was found to interact with 26 MPGs overall, 22 are highly significant. Regarding the highly interacting MPGs, three showed the most interactions with our coexpression modules: formaldehyde biosynthesis from methylamine (PWY-6965), L-lysine degradation to glutaryl-CoA (PWY-6328) and putrescine degradation to 4-aminobutanoate (PUTDEG-PWY). Altogether, the bipartite network indicated the environmentally sensitive modules as well as their potential triggers.

**Figure 7.**
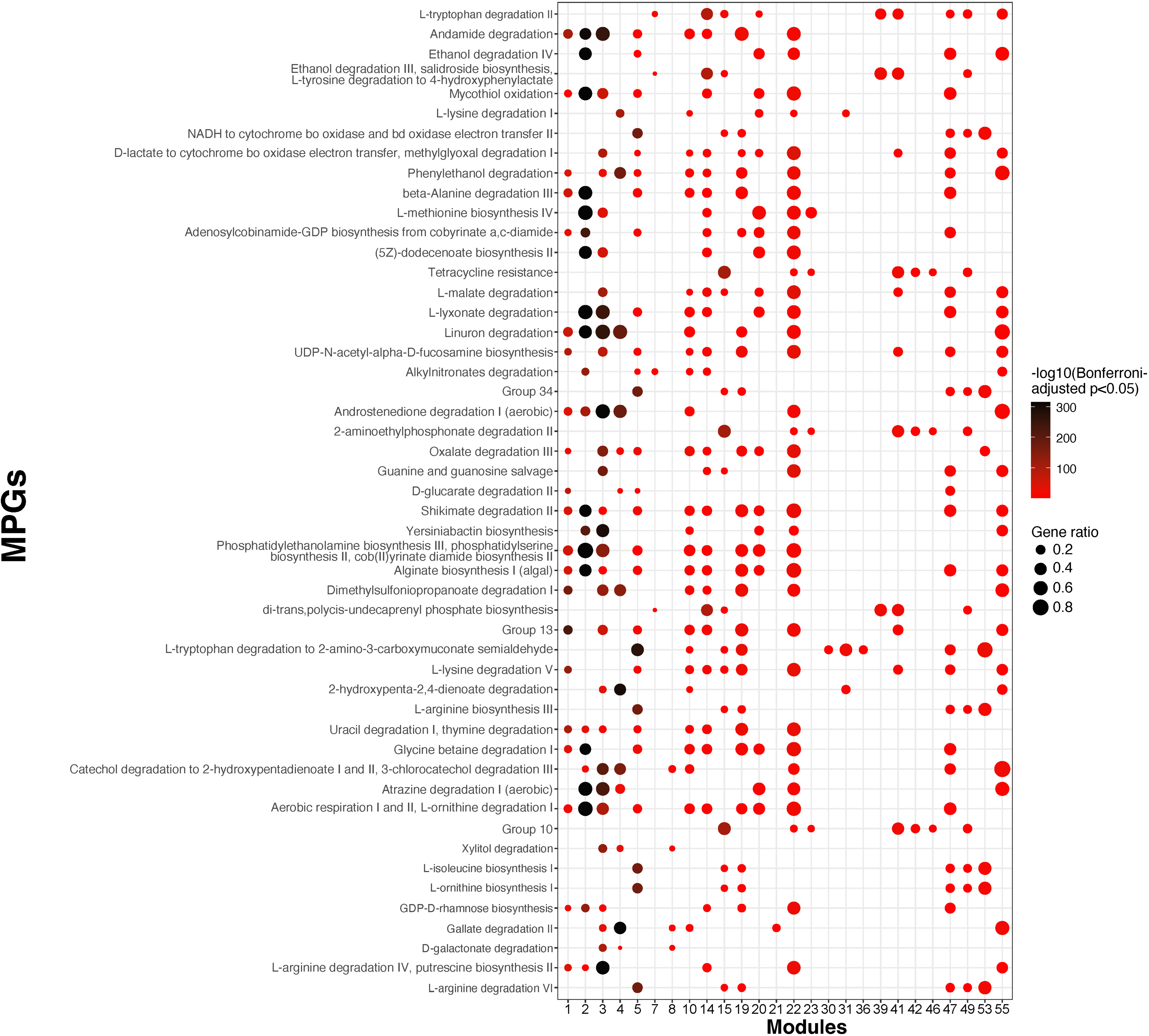
Interactions between bacterial pathways and coexpression modules. (A) A bipartite network was built for MPGs and coexpression modules by defining interactions based on the significance of the gene set overlap (lowest Bonferroni-adjusted p-value<0.05). The plot shows all interactions between selected MPGs that were defined based on the 50 most significant interactions and the top 60 modules. The p-value scale depicts the negative logarithm of the corrected p-values.

## Discussion

All animals live in tight associations with microbial communities and bacteria can have a significant impact on the biology of their host. In nematodes, bacteria have been demonstrated to affect several traits including development, pathogen resistance, and behavior^2, 5, 20, 39, 40^. After observing such intriguing phenotypes, the key question arises what bacterial signal triggers this response. While one major advantage of the nematode system is that individual interactions can be studied in isolation, we combine data from many different bacteria to ask broader questions such as to what extent does the nematode’s transcriptomic response vary across bacterial diets. Can these responses be traced back to specific metabolic pathways in the bacteria? These systems level questions distinguish our work from many other studies that focused on specific effects of individual bacteria^2, 5, 20, 39, 40^. By investigating a large number of bacterial diets, we aimed to have higher statistical power to also detect a greater number of metabolic interactions between dietary bacteria and *P. pacificus* nematodes. It should be noted that the capability to detect metabolic interaction will depend on the sampling of bacteria and the phylogenetic distribution of the pathway. For this reason, we have tried to maximize the number of sequenced samples, which resulted in a collection of 84 bacterial genomes that serve as a genomic basis for the current and future research on host-microbe interactions. At the same time, this data allowed us to investigate the extent of variation in metabolic potential within and between bacterial genera, which revealed considerable amounts of intra-genus level variation in bacterial metabolic potential as well as in the nematode transcriptomic response. Following a similar approach as Zimmermann et al.^41^, who associated metabolic potential of members of the *C. elegans* microbiome with specific traits, we made use of previously generated phenotypic data to perform an inter-species association study to identify potential connections between bacterial pathways and nematode behavior and survival. In particular, we could confirm the nematicidal activity of a certain coumarin-derivative on *P. pacificus*.

The generation of RNA-seq data of *P. pacificus* nematodes for 38 different bacterial diets represents to our knowledge the largest transcriptomic study in the field of interactions between nematodes and bacteria. This transcriptomic data placed us in a unique position to perform a large-scale interspecies association study, where we tested the absence/presence pattern of an MPG for a significant expression response of a given coexpression module. This identified more than 2,800 associations between bacterial metabolic pathways and specific coexpression modules. Each of these individual interactions implies a causal relationship between one or several metabolites in a bacterial metabolic pathway and the response of a specific coexpression module in the worm. The shear number of associations points towards a highly complex network of metabolic interactions between *P. pacificus* and its dietary bacteria. However, the actual number might be smaller, as similar absence/presence patterns among MPGs will likely predict the same interacting coexpression modules. At the same time, the number of interactions will be underestimated due to the lack of statistical power to detect interactions for MPGs that show very little variation among the bacterial genomes. Another uncertainty comes from the fact that some of the transcriptomic responses may represent indirect effects of altered development. Thus, future studies could employ additional transcriptomic data at a much higher temporal resolution to dissect immediate responses in response to bacteria from indirect effects that are mediated through altered development. This will simplify the selection of candidate metabolites for supplementation experiments in order to validate predicted interactions. Knowing how different metabolites affect the transcriptional state of the worm, will ultimately allow us to manipulate the system in a controlled manner.

## Methods

### Key resources table

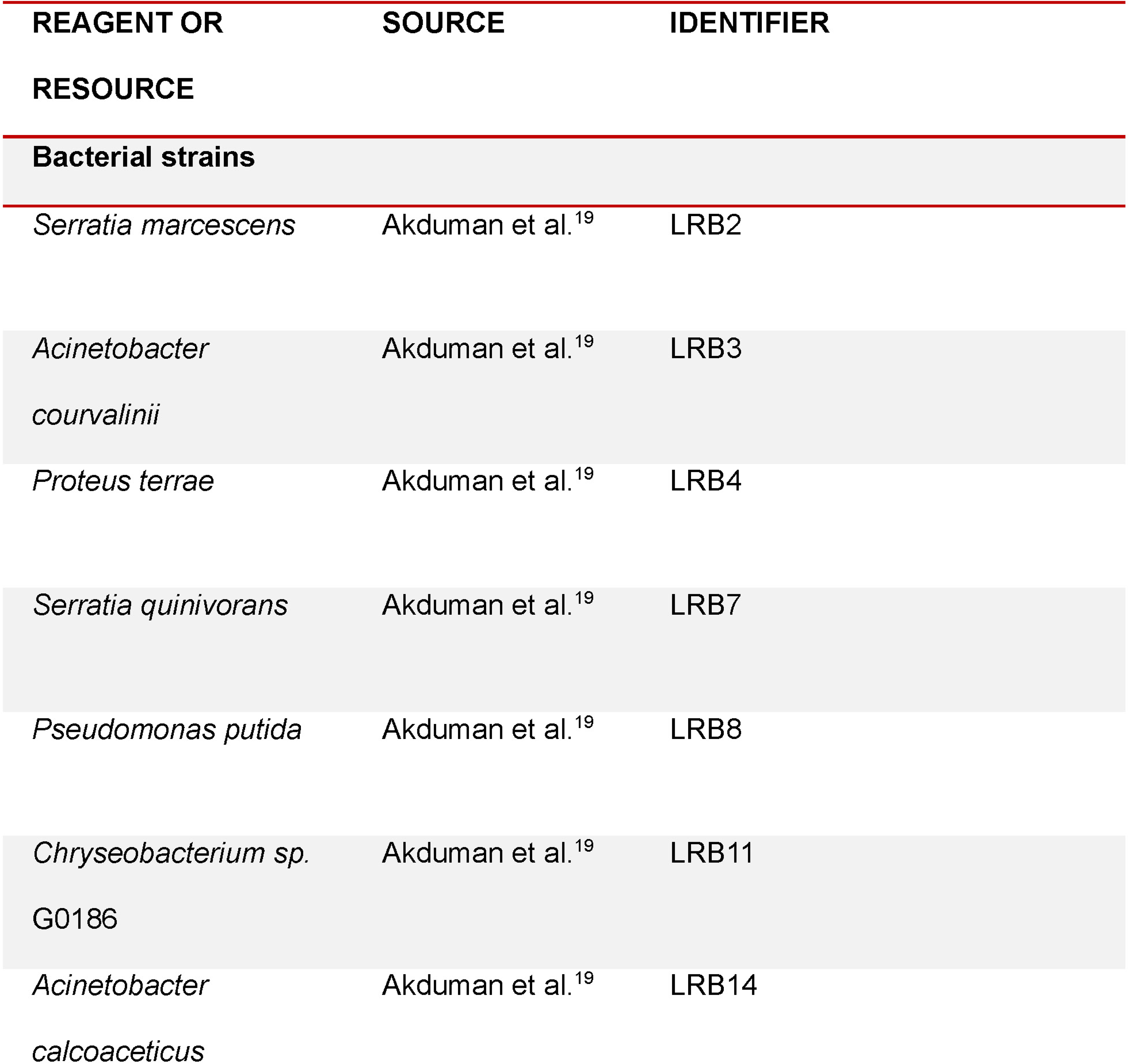
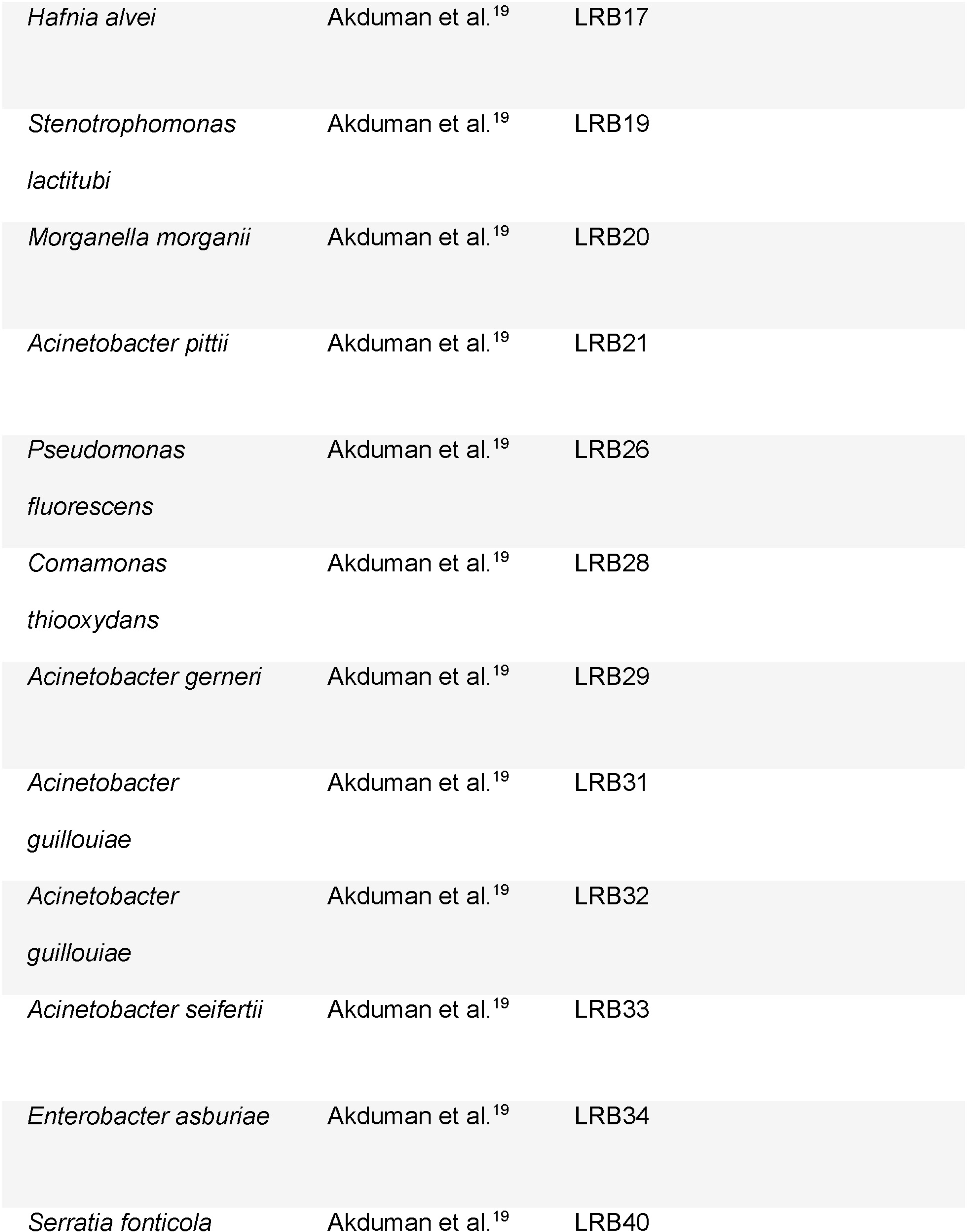
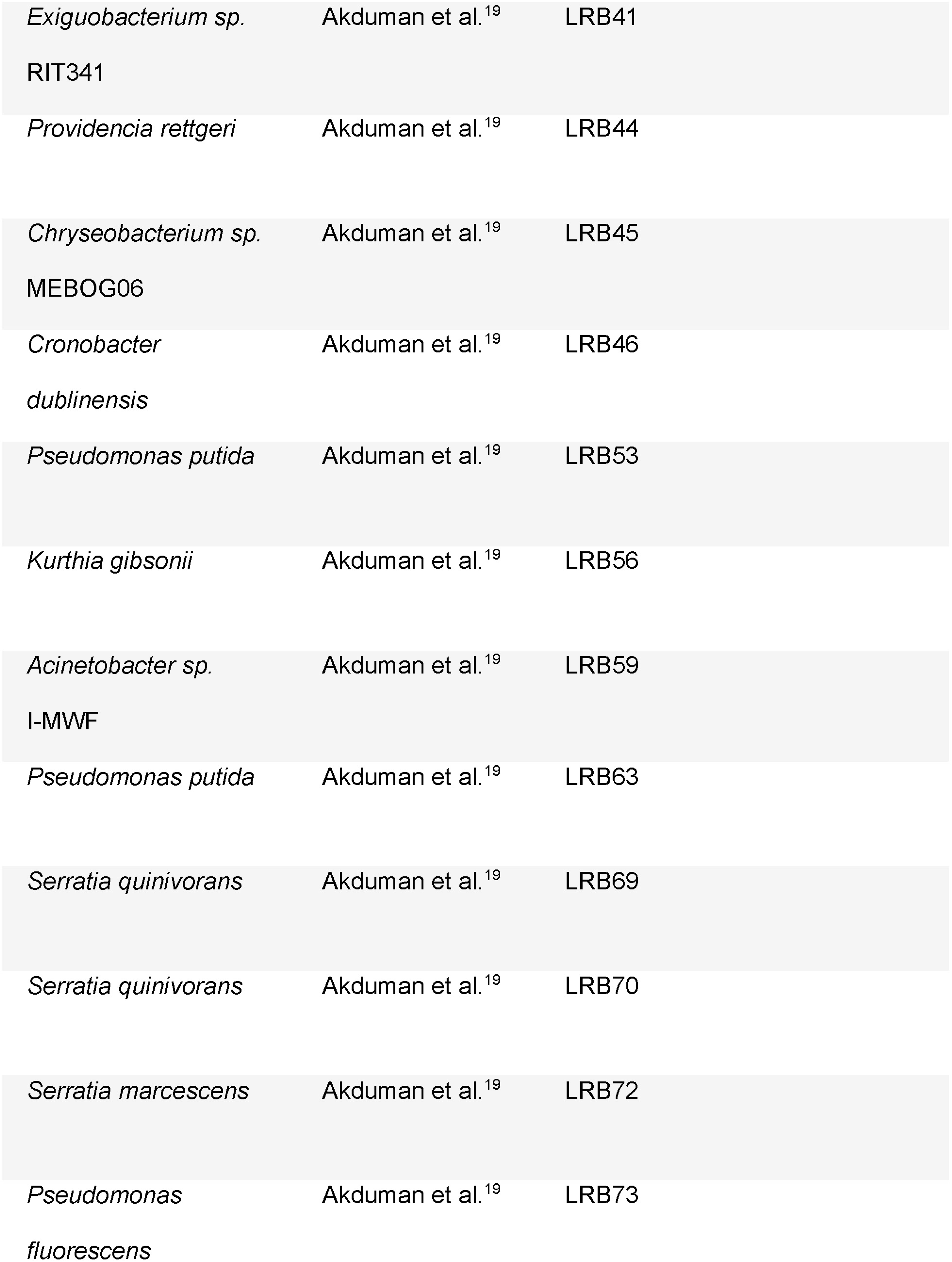
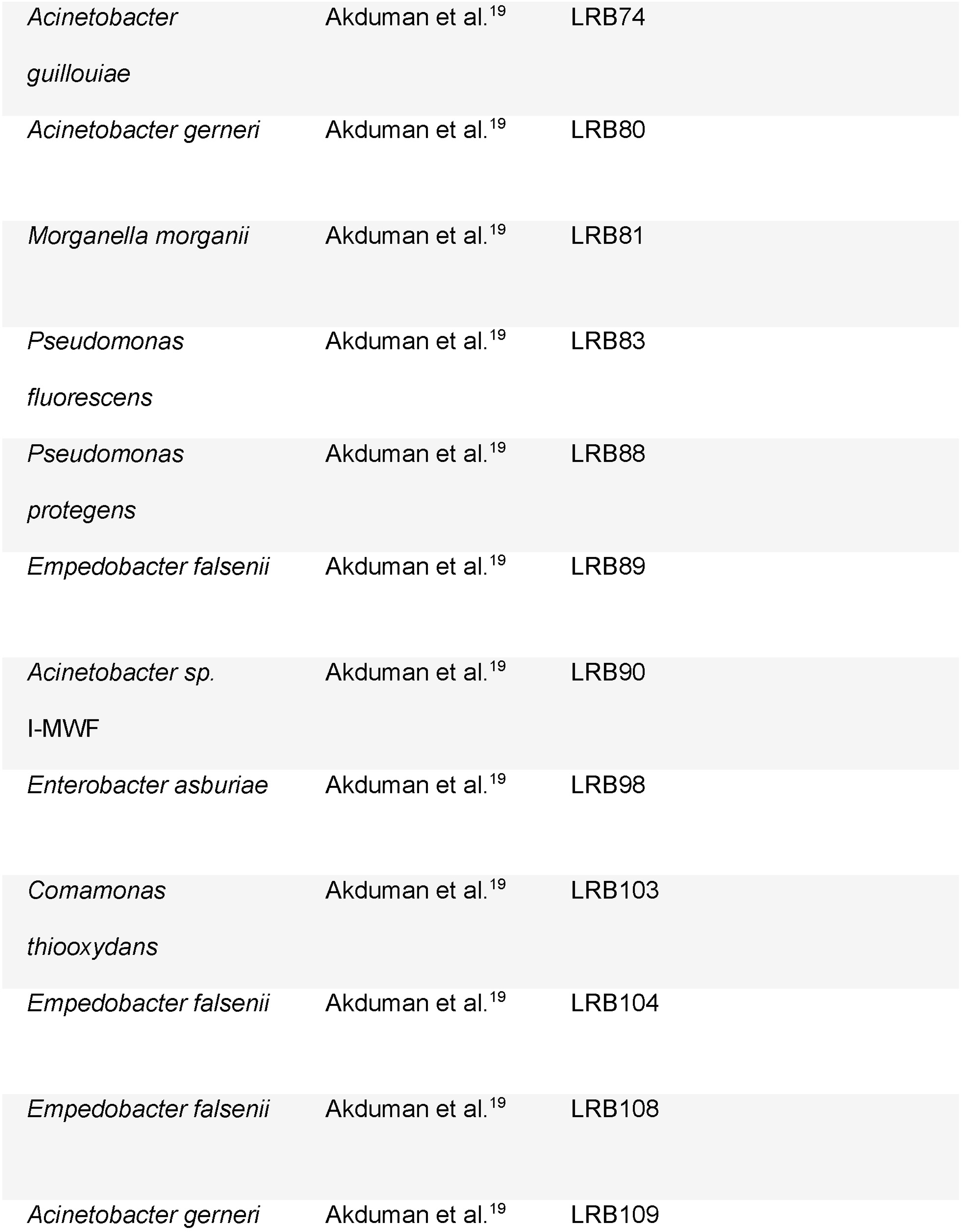
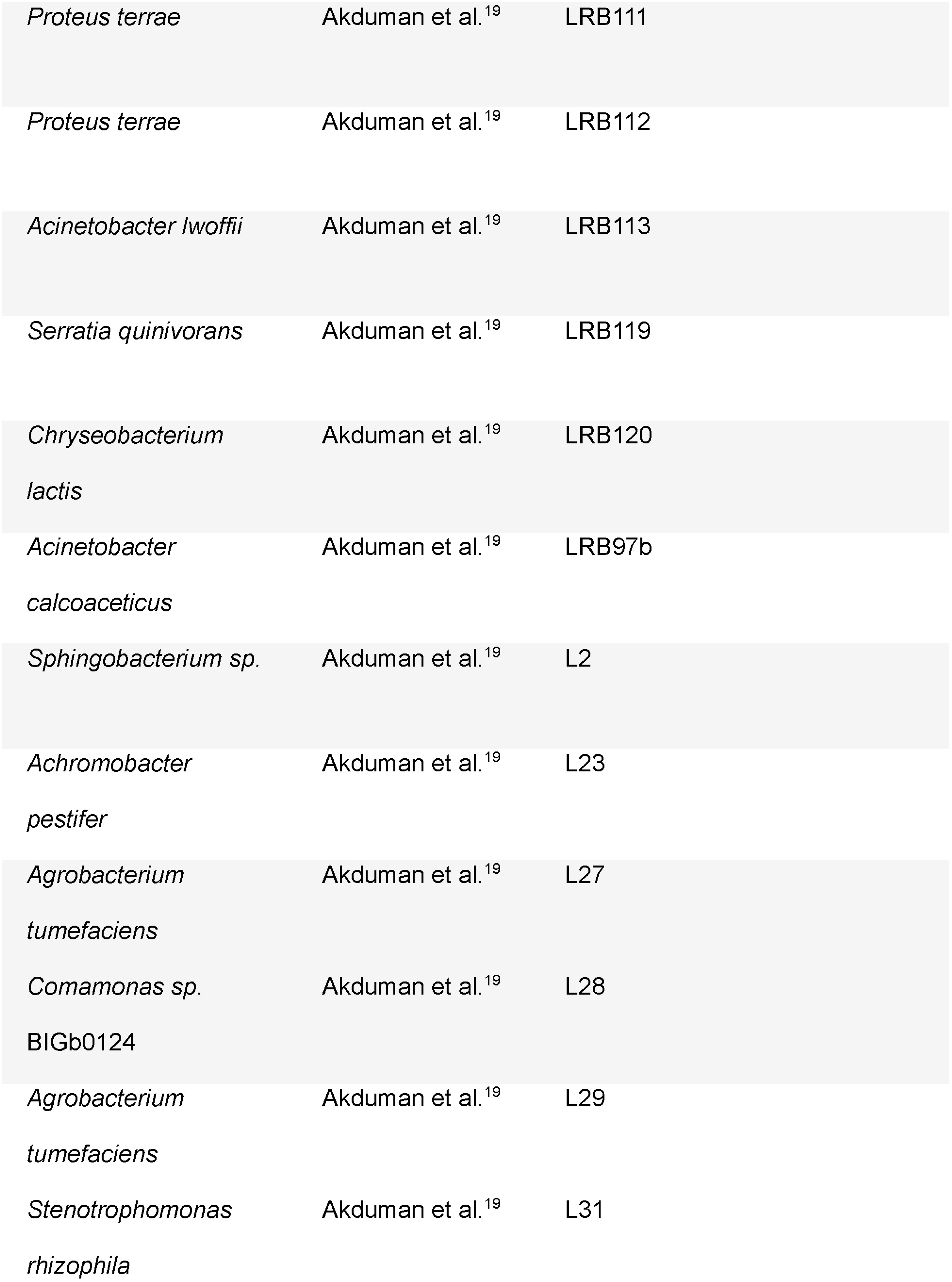
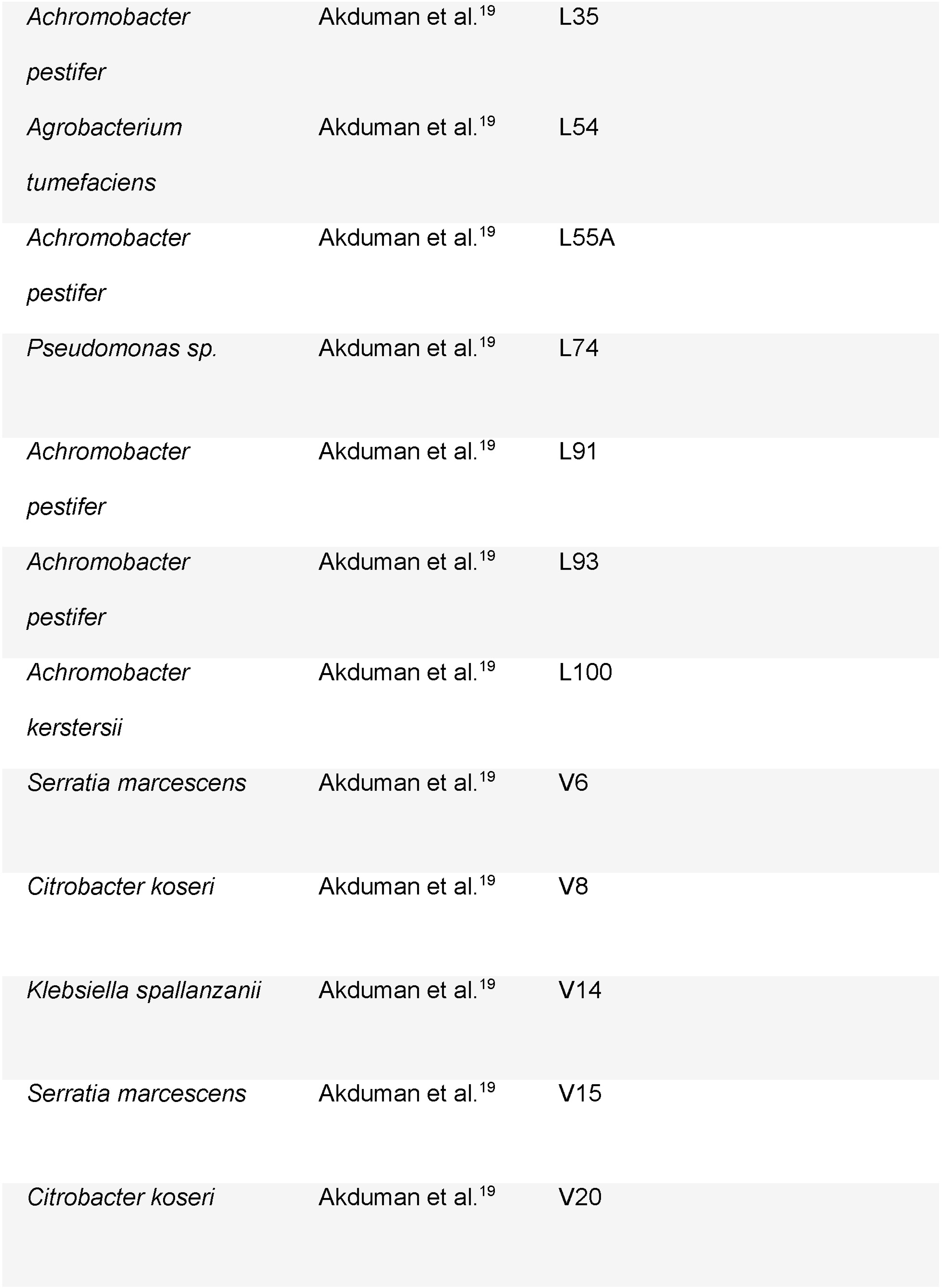
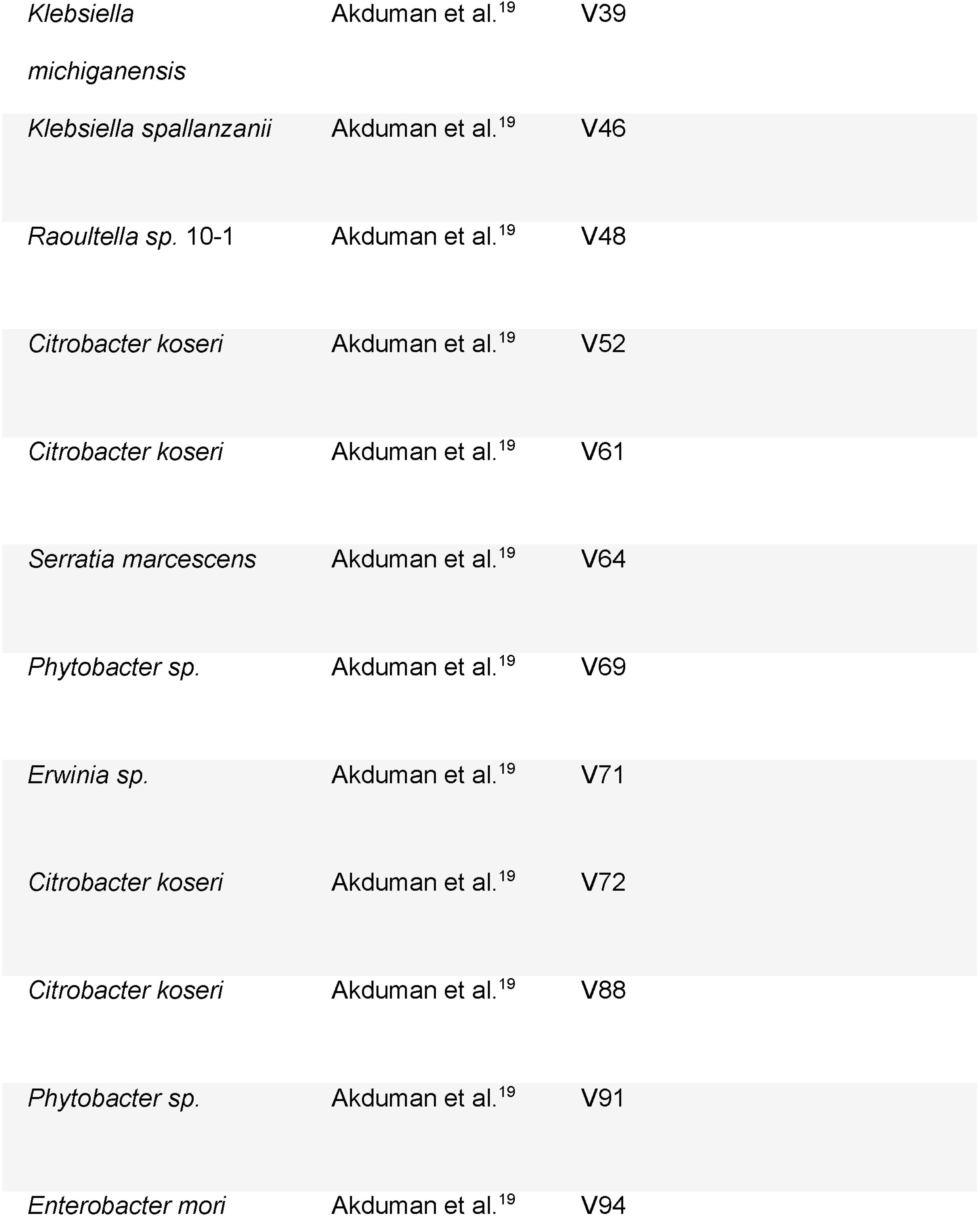
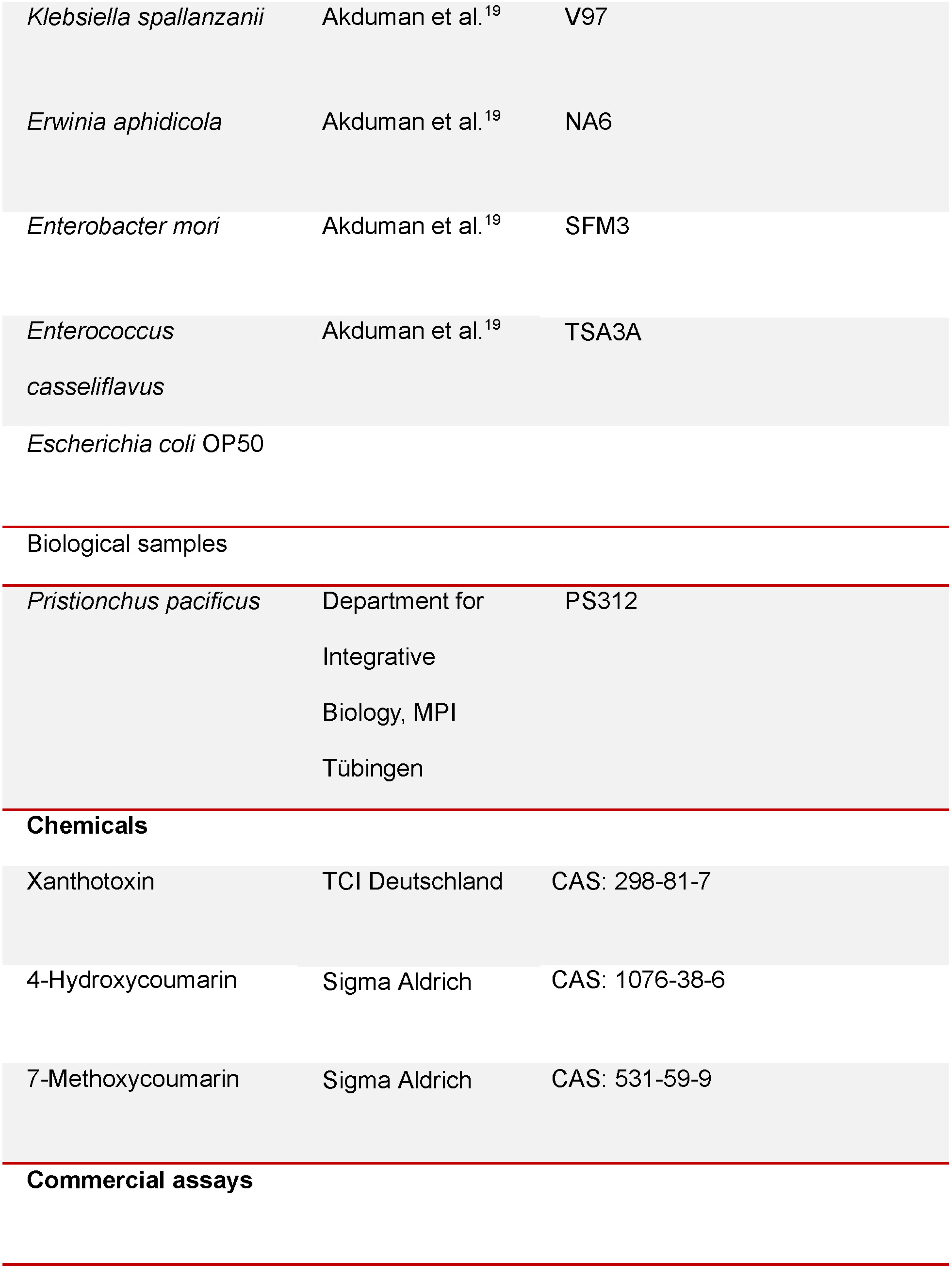
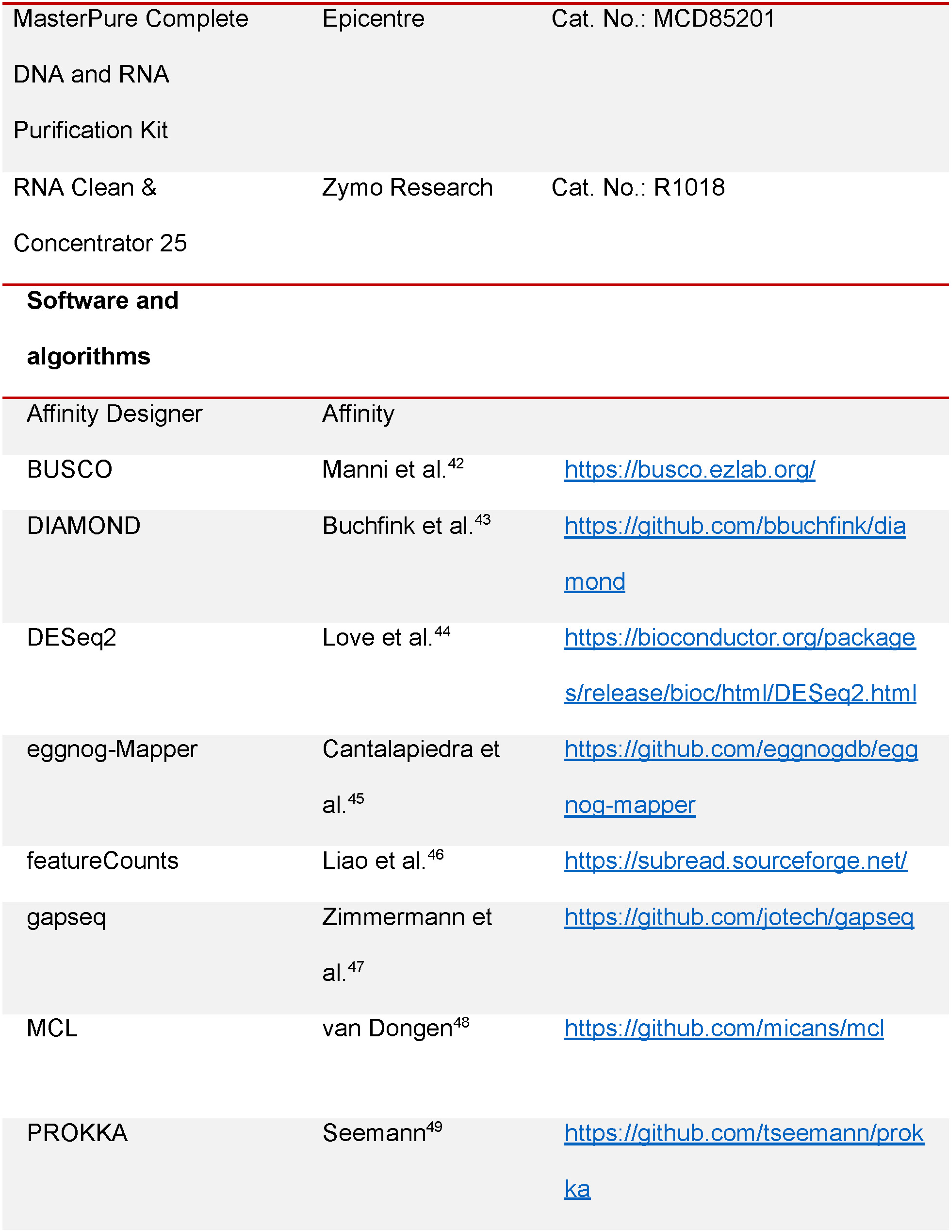
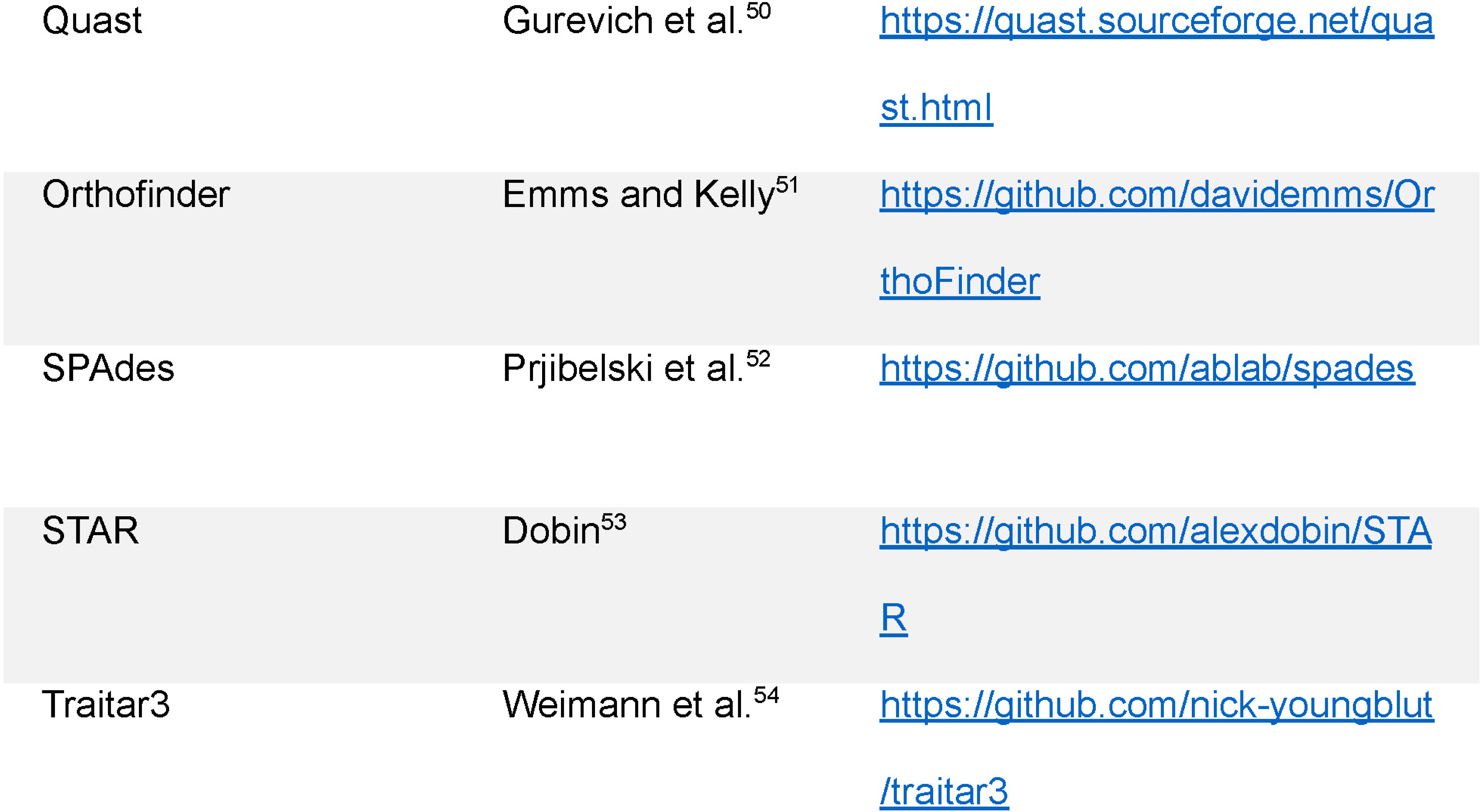

### Resource availability

#### Materials availability

This study did not generate new unique resources and reagents.

#### Data and code availability

The raw sequencing data for the bacterial genomes, the genome assemblies and the raw read counts of the transcriptomic profiles from this study have been deposited at the European Nucleotide Archive under Bioproject PRJEB80633.

### Experimental model details

#### Bacterial culture conditions

All bacterial strains were initially restreaked from a glycerol stock on Lysogeny Broth (LB) plates and grown overnight in LB. Bacteria were grown at 30 °C or 37 °C depending on the species.

#### Nematode growth conditions

The wild type strain of *Pristionchus pacificus* (PS312) was maintained at 20 °C on nematode growth medium (NGM) seeded with *Escherichia coli* OP50 before use. From every generation, three adults were transferred to fresh NGM plates with a worm pick.

### Method details

#### Whole genome sequencing of bacterial strains

All bacterial strains to be sequenced were grown overnight in LB in 15 ml falcon tubes and DNA samples were obtained using the Epicentre MasterPure Complete DNA and RNA Purification Kit (Illumina, San Diego, USA). DNA libraries were prepared using the Illumina DNA Prep kit according to the manufacturer’s guidelines. The libraries were quantified using both a Qubit 2.0 Fluorometer (Thermo Fisher Scientific, Waltham, USA) and a Bioanalyzer (Agilent Technologies, Santa Clara, California) and normalized to 2.5 nM. Samples were sequenced as 150bp single-end reads on multiplexed lanes of an Illumina HiSeq3000 in our in-house sequencing facility. Raw reads were deposited at the European Nucleotide Archive (ENA) under the study accession PRJEB80633.

#### Genome assembly and metabolic potential identification of bacterial strains

The bacterial genomes were assembled from the raw reads using SPAdes^52^ (version 3.15.1, parameters: --careful -o -1 -2 -t 30 –cov-cutoff ‘auto’) and annotated with PROKKA^49^ (version 1.14.6, parameters: --addgenes) and eggNOG-Mapper^45^ (version 2.1.0-1, eggnog DB version: 5.0.2). The predicted bacterial protein sequences were used as input to detect orthogroups with OrthoFinder and reconstruct the phylogeny with the incorporated RAxML^51^. The bacterial genomes were used to predict the presence or absence of metabolic pathways with gapseq^47^ (version 1.2, parameters: find -p all, MetaCyc^55^ as the default database) as well as the presence of 67 bacterial phenotypes with the implementation of Traitar in python (traitar3^54^, version 3.0.1, parameters: predict). The bacterial metabolic pathways were grouped (MPGs) based on their presence/absence patterns across the bacterial strains to reduce redundancy. This decreased the number of bacterial metabolic pathways to be assessed from 2902 to 712. The nomenclature of the bacterial strains was determined by applying DIAMOND^43^ (version 2.1.4.158, parameters: makedb and blastp) to compare the predicted bacterial protein sequences against the non-redundant version of the NCBI database (August 2023 version).

#### Association of MPGs with chemotaxis and survival

In the initial study, the chemotaxis index ranges from -1 (repulsion) to 1 (attraction) and the survival index from 0 to 100, the latter reflecting the percentage of worms that were alive on the 5^th^ day post transfer^19^. For our analysis, we binarized the assay data by assigning the values below 40 percent survival and 0.2 chemotaxis as zero and the rest as one. The threshold for the binarization was determined based on the distribution of values. To test the association between each MPG and the nematode phenotypes, we divided the assay data to two vectors, representing the set of values when the nematode was maintained on bacterial strains with and without the MPG and performed a Mann-Whitney-U test, followed by Bonferroni-correction (P < 0.05). To associate bacterial virulence factors with survival, we downloaded the 4,236 core proteins from the Virulence Factor DataBase^30^ (2024-07-31) and performed pairwise all-against-all BLASTP searches for all bacterial protein sets combined with the VFDB data (e-value < 10^-5^). The widely used clustering algorithm mcl was used to cluster the data into 1605 orthogroups. These orthogroups were then tested for association with survival using a Wilcoxon-test (FDR-corrected P < 0.1).

#### Supplementation experiments

To check the nematicidal effect of coumarin derivatives *in vivo*, we supplemented 4-Hydroxycoumarin, 7-Methoxycoumarin and xanthotoxin to the nematode. The coumarins were added directly to liquid NGM to a final concentration of 1mM and the solution was subsequently poured to 6 cm plates (11 ml per plate). The plates were seeded with 300 μl of *E. coli* OP50 the next day and left to grow a bacterial lawn for two days. After that time, 20 young adult hermaphrodites were transferred on each plate and their survival was tracked daily for five days. The surviving nematodes were transferred on new plates containing the coumarins on the third day after the initial transfer to avoid food depletion and misidentification from their offspring. Mortality was determined by lack of response to prodding with the pick. For the control and each coumarin, we performed five replicates.

#### Dietary experiments

Eggs from *P. pacificus* adults were obtained and spotted on NGM plates seeded with 75 μl bacterial overnight cultures to produce the parental generation (P0) on each diet. 25 gravid hermaphrodites from the P0 were transferred to a new plate seeded with the same bacterial strain and were left to lay eggs for five hours, after which point they were removed. The F1 worms were collected 72 hours after the initial transfer. Collection of the worms required washing the plate with autoclaved M9^-^ buffer, centrifugation at 500g for a minute and discarding the supernatant in order to remove as much of the collected bacterial lawn as possible. The worm pellets were flash frozen in liquid N_2_ and were used for RNA extraction or stored at -80°C for later processing.

#### RNA sequencing

For total RNA extraction, the frozen worm pellets were treated with Trizole followed by purification with the Zymo RNA Clean & Concentrator 25 Kit according to the manufacturer’s instructions. The extracted RNA was quantified and quality assessed with a NanoDrop ND1000 spectrometer (PeqLab, Erlangen, Germany) and a Qubit 2.0 Fluorometer. The samples were shipped to Novogene for library preparation and messenger RNA (mRNA) sequencing. Libraries were sequenced as 150bp paired-end reads on an Illumina NovoSeq 6000 platform. The obtained raw reads have been deposited in the European Nucleotide Archive under the study accession number PRJEB80633.

#### Gene coexpression network

The raw reads were aligned to the reference *Pristionchus pacificus* genome with STAR^53^ (version 2.7.1a) and quantified with featureCounts^46^ from the Subread R package (version 2.0.1) based on the latest annotations. By filtering the count matrix to remove genes with less than 10 reads total, we reduced the number of *P. pacificus* genes to be assessed from 28896 to 24335. The read counts across different conditions and replicates were normalized with the DESeq2^44^ counts function (option: normalized = TRUE) and were input in MCL^48^ (version 22-282, r=0.7, I=2) to create a gene coexpression network. The network modules were tested for the enrichment of protein domain annotations (PFAM)^33^, metabolic pathways (KEGG)^35^ and gene expression sets from previous works^25^ with Fisher’s exact-test followed by Bonferroni correction (P < 0.05) in R (version 4.4.0). The network modules were assessed further for their interactions with the bacterial metabolic potential. Specifically, for each MPG, the normalized counts of the genes in each module were separated into two vectors based on whether the presence or absence of the MPG in the bacterial strain in order to represent the expression of a specific gene in a module in the presence or absence of a MPG. The presence and absence expression vectors for each gene were compared using Mann-Whitney-U test as well as to calculate the fold change in expression. To determine the interaction of a module with a MPG, we retained the genes where the fold change was lower than 0.5 or higher than 2 and performed multiple hypothesis testing for the module genes that fulfilled the condition (Bonferroni < 0.1). The list of genes identified as interacting with the MPGs were tested for module enrichment using Fisher’s exact test (Bonferroni-adjusted p-value<0.05) to determine the interaction between the MPGs and the coexpression modules. The bipartite network was implemented using Python (version 3.10.12).

## Notes

### Competing Interest Statement

The authors have declared no competing interest.

