## Supplemental figures for "Interspecies systems biology links bacterial metabolic pathways to nematode gene expression, behavior, and survival"

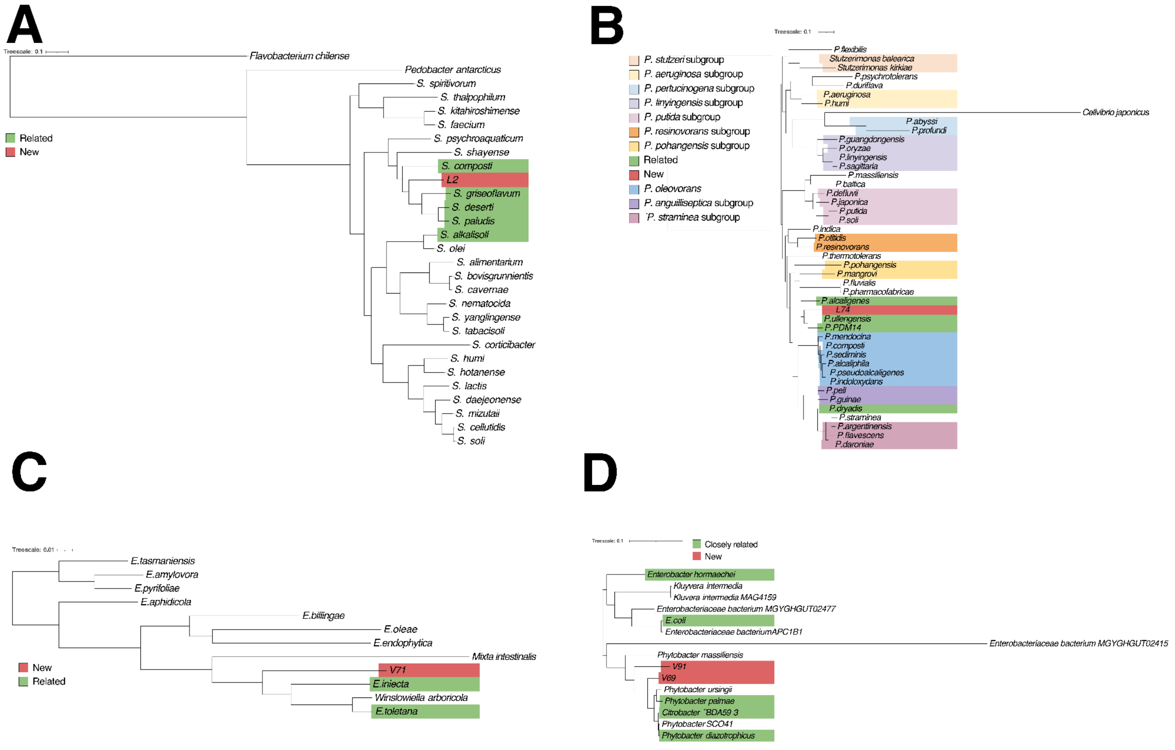


**Supplemental Figure S1. Reconstruction of bacterial phylogenies to characterize novel strains.** Potentially novel bacterial strains in our collection were integrated into previously published phylogenies by reconstructing species trees that include the candidate genome together with their best hits in the non-redundant version of NCBI. (A) The phylogeny for *Sphingobacterium* L2 (B) The phylogeny for *Pseudomonas* L74. (C) The phylogeny for *Erwinia* V71. (D) The phylogeny for *Phytobacter* V69 and V91.

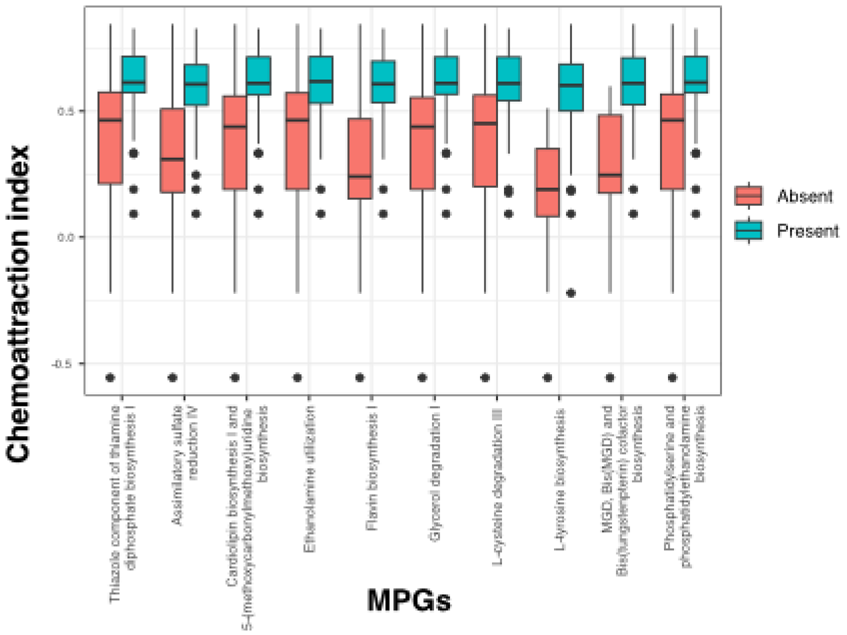


**Supplemental Figure S2. Association of MPGs with chemoattraction in the nematode.** Association analysis with the MPGs against the chemoattraction data from Akduman et al (2018) identified 25 MPGs with a predicted significant impact on chemotaxis behavior, a subset of which is depicted here.

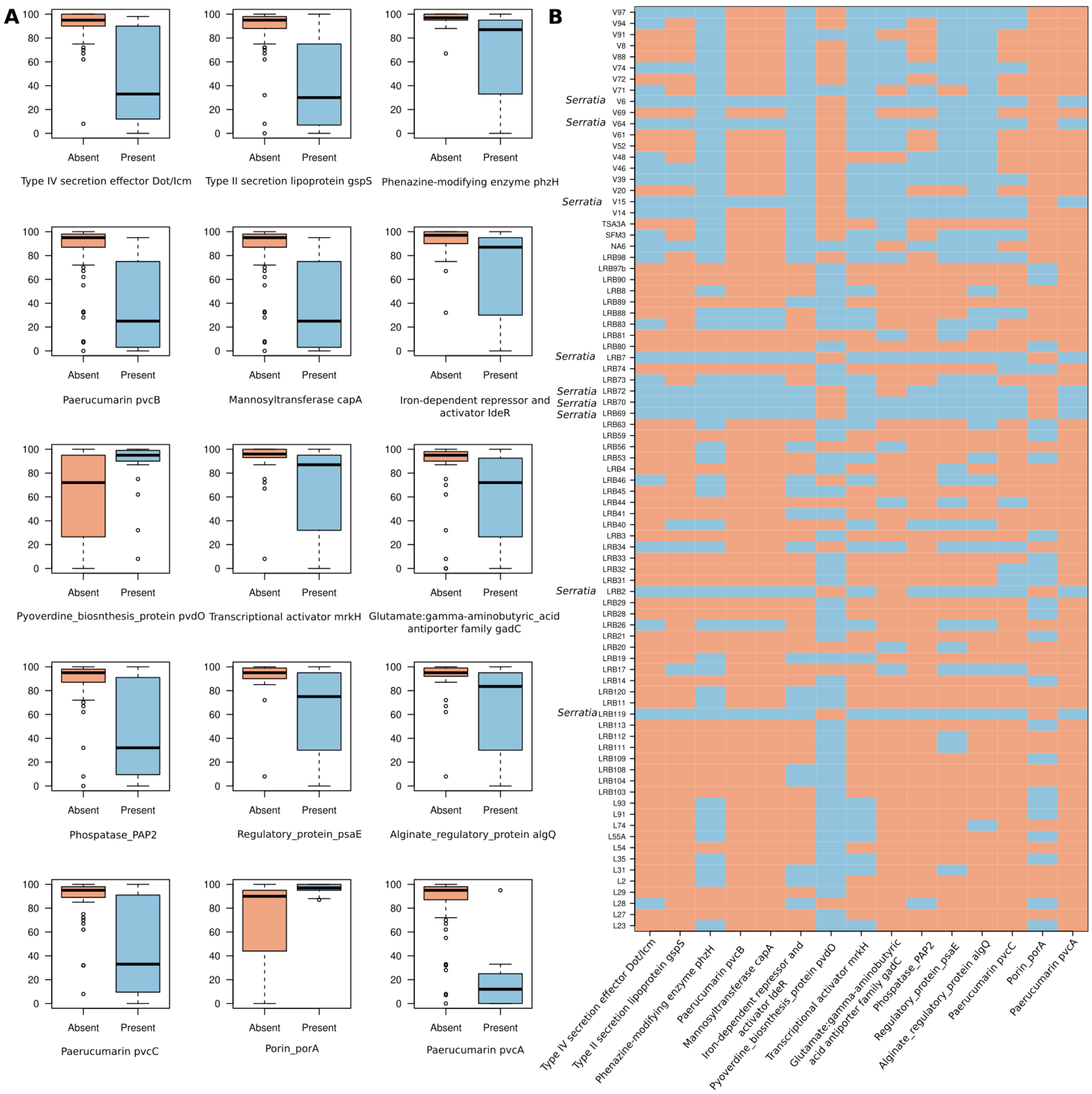


**Supplemental Figure S3. Association of bacterial virulence factors with nematode survival.** A) The core protein set of the Virulence Factor DataBase (VFDB) analysis was clustered together with the bacterial protein sets into orthogroups which were then used to test for associations with survival data. The boxplots show the most significantly associated orthogroups, labeled with a representative VFDB member. B) The heatmap shows the presence/absence pattern of the candidate orthogroups across the bacterial genomes.

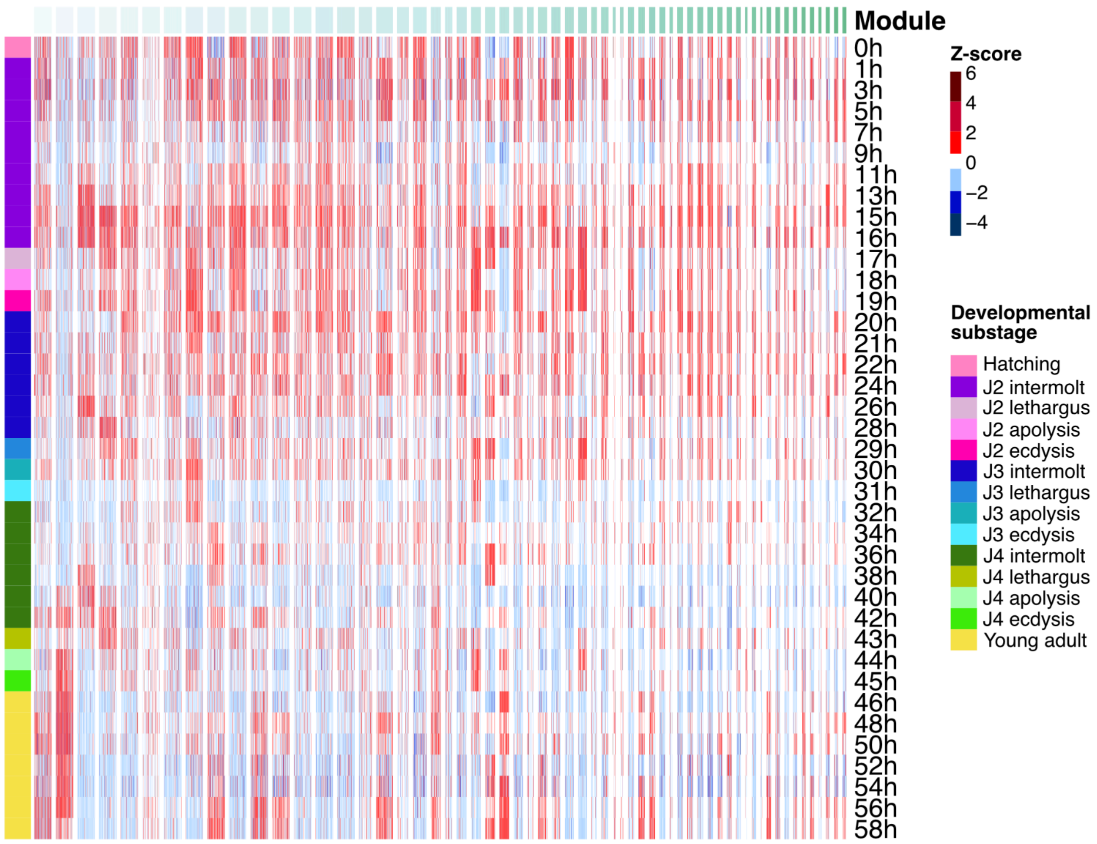


**Supplemental Figure S4. Visualization of the co-expression modules with developmental time course data from Sun et al (2021).** The top modules of the co-expression network were visualized with developmental time-course data. This revealed developmental signatures in various modules.
